# Joint signatures of morphological and microstructural inter-individual variation in the Alzheimer’s spectrum

**DOI:** 10.1101/2024.01.24.576996

**Authors:** Aurélie Bussy, Raihaan Patel, Olivier Parent, Alyssa Salaciak, Saashi A. Bedford, Sarah Farzin, Stephanie Tullo, Cynthia Picard, Sylvia Villeneuve, Judes Poirier, John CS Breitner, Gabriel A. Devenyi, PREVENT-AD Research Group, Christine L. Tardif, M. Mallar Chakravarty

## Abstract

Alzheimer’s disease (AD) is primarily characterized by the accumulation of amyloid and tau pathologies. However, alterations in the detailed organization and composition of neural tissue also contribute to the disease’s early stages. Here, we sought to explore whether hippocampal and cortical microstructural changes, such as myelin alterations and inflammation-mediated increases in iron, could serve as indices of AD-related pathophysiology. In this study, we included 158 participants across the AD spectrum: from individuals without cognitive impairment, at high risk for AD, in the prodromal phase with mild cognitive impairment, and suffering from clinical dementia. We measured atrophy using structural magnetic resonance imaging (MRI) and estimated myelin and iron content using quantitative MRI (qMRI) metrics derived from T1 and T2* relaxation, times respectively. We integrated these contrasts to estimate a joint multivariate signature of tissue alterations across the cortex and hippocampus using non-negative matrix factorization. The relevance of these signatures to AD-spectrum measures of medical history, lifestyle, and cognition were further explored using partial least squares correlation. Our results reveal lower disease-related cortical thickness over large areas of the cortex while T2* provided specific variation across the brain (lower in dorsomedial and superior temporal areas, superior frontal cortex, and premotor cortex, and higher in the occipital lobe). Additionally, we observed longer T1 and T2* times in the hippocampus associated with specific lifestyle risk factors like past smoking, high blood pressure, high cholesterol levels, and higher anxiety. These patterns were significantly related to older age, associated with AD progression, being female, and being an APOE-□4 carrier. Taken together, our results suggest that qMRI metrics could serve as a valuable non-invasive tool for exploring the role of myelin and inflammation in AD-related pathophysiology and could be sensitive to modifiable risk factors related to lifestyle and medical history. Future studies may use these signatures to investigate their relationship in investigations related to lifestyle interventions or novel therapeutics.

## 1. Introduction

As the global population ages, the proportion of elderly individuals is increasing at an unprecedented rate. Advanced age is the most significant risk factor for neurodegenerative diseases, such as Alzheimer’s disease (AD)^1^. The recently approved AD-related therapies^2,3^ mostly target extracellular deposition of ß-amyloid (Aß) with modest effect sizes on outcome measures that are important to patients: namely cognition and activities of daily living^4^. Further, while Aß and intracellular accumulation of hyperphosphorylated microtubule associated protein tau (p-tau) are the main pathological hallmarks of AD, the disease is known to be multifactorial and to involve other microstructural changes such as myelin deterioration and inflammation-mediated increases in iron^5^. Additionally, the above pathological processes are likely initiated several years if not decades before the first clinical manifestations of the disease^6^. Here, we propose to examine microstructural and morphological signatures of the disease across the AD spectrum: from individuals not suffering from cognitive impairment, those at high risk for AD, those in the prodromal phase with mild cognitive impairment (MCI), and those suffering from clinical dementia.

To assess the multifactorial aspect of AD progression, we used a combination of neuroimaging measures that index both local brain atrophy and microstructure. Specifically, quantitative MRI (qMRI) techniques were used to estimate the biophysical properties of the underlying tissue sample (i.e amount of myelin and iron) through the use of relaxation times. While qMRI has been increasingly used in some clinical research related to multiple sclerosis^7^ or musculoskeletal diseases^8^, it has been under-explored in the context of the AD spectrum^9^. Nonetheless, it has been demonstrated that myelin breakdown and inflammation-induced iron accumulation may promote amyloid accumulation^10–14^ and that individuals in the earliest stages of AD harbour a higher iron load^15–17^. These previous findings strongly suggest that myelin and iron are two pathologically-relevant microstructural properties in the context of AD, and their *in vivo* measurement could add neurobiological insights regarding disease pathophysiology and complement treatment targets.

In this study, we used a data integration technique, namely non-negative matrix factorization (NMF), to derive components defined by groupings of voxels sharing common modes of covariation across MRI-based measures^18^. This strategy is ideal to identify relevant brain regions while simultaneously providing insight into biological sensitivity and specificity within these regions. For example, morphological measures lack a precise biological interpretation, and incorporation of microstructural information from qMRI can provide insights into the putative underlying causes of volume changes. Here, we specifically studied the hippocampus^19,20^ and cortex^21–23^ because structural MRI studies have repeatedly identified that morphometric changes in these areas are reliable indicators of AD pathology. To better understand the association between NMF-derived brain patterns and relevant factors in the AD-spectrum, we examined their multi-variate relationship to demographics, medical, cognitive information and lifestyle risk factors.

Our findings suggest that the cortical morphometry and hippocampal microstructure patterns may sensitively index AD progression. Specifically, our microstructural findings reveal lower tissue integrity, decreased myelin, higher water content, and iron reduction in the hippocampus, all of which are associated with disease stage. Cortical findings revealed a reduction of cortical thickness (CT) and surface area (SA), but no myelin changes, suggesting a neuronal loss more than a glial alteration related to the progression of AD. Finally, multivariate analyses demonstrate an association between some of these patterns that were preferentially associated with AD-related risk factors like smoking, high cholesterol levels, blood pressure, and anxiety. We believe that this approach can be further refined to examine the impact of trials related to lifestyle interventions or novel therapeutics.

## 2. Methods

A workflow of the key methodological steps employed in our study is depicted in **Figure 1**.

**Figure 1:**
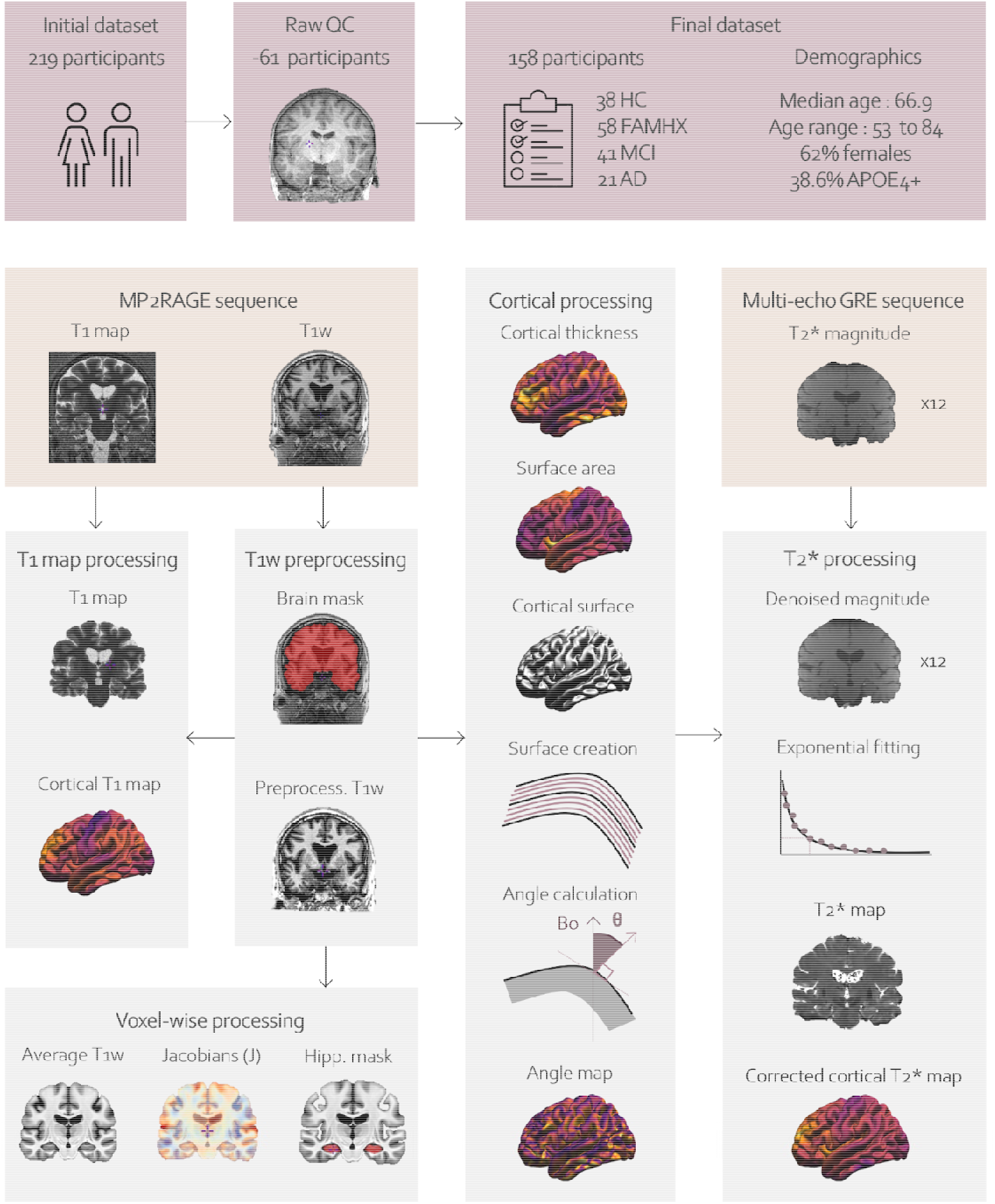
Schematic representation of the methods. From the 219 initial participants, 61 were excluded due to motion quality control (QC), leaving 158 participants with high-quality MRI images that were included in the final sample. The MP2RAGE sequence provides a T1w image and a quantitative T1 map. The minc-bpipe-library is used to preprocess the T1w images and obtain a brain mask. CIVET is used to extract CT, SA and cortical surfaces. Additional surfaces were created to sample the maps at different cortical depths (12.5%, 25%, 37.5%, 50%, 62.5%, 75% and 87.5%). □ is defined as the angle between □*_0_* and the cortical normal at each vertex. The multi-echo gradient echo sequence provides 12 magnitude images. A denoising using adaptive non-local means denoising ^24^ is applied on each echo. An exponential fit was used to extract T2* relaxation times from the 12 echoes and the angle □ was used to residualized the cortical T2* values. Deformation based morphometry (DBM) was used to calculate jacobians (J) using the T1w scans. Using the average template created from DBM, we manually defined a hippocampal mask to extract hippocampal voxel-wise metrics.

### 2.1. Participants

Individuals from two datasets collected at the Douglas Research Center were included. This data acquisition was approved by McGill Institutional Review Board and Douglas Mental Health University Institute Research Ethics Board. 219 participants were originally included: 168 individuals were part of the Alzheimer’s disease biomarker cohort^25–29^ and 51 were part of the “Pre-symptomatic Evaluation of Experimental or Novel Treatments for Alzheimer Disease” (PREVENT-AD) cohort^30^. Inclusion and exclusion criteria are described in Supplementary methods. Demographics of our participants are reported in **Table 1**. After quality control (QC) of the image processing (described in **Sections 2.2, 2.4 and 2.6**), we included 158 individuals across the AD spectrum: 38 healthy controls (HC), 58 cognitively healthy individuals with a parental AD history (FAMHX), 41 with mild cognitive impairment (MCI) and 21 with AD.

**Table 1:**
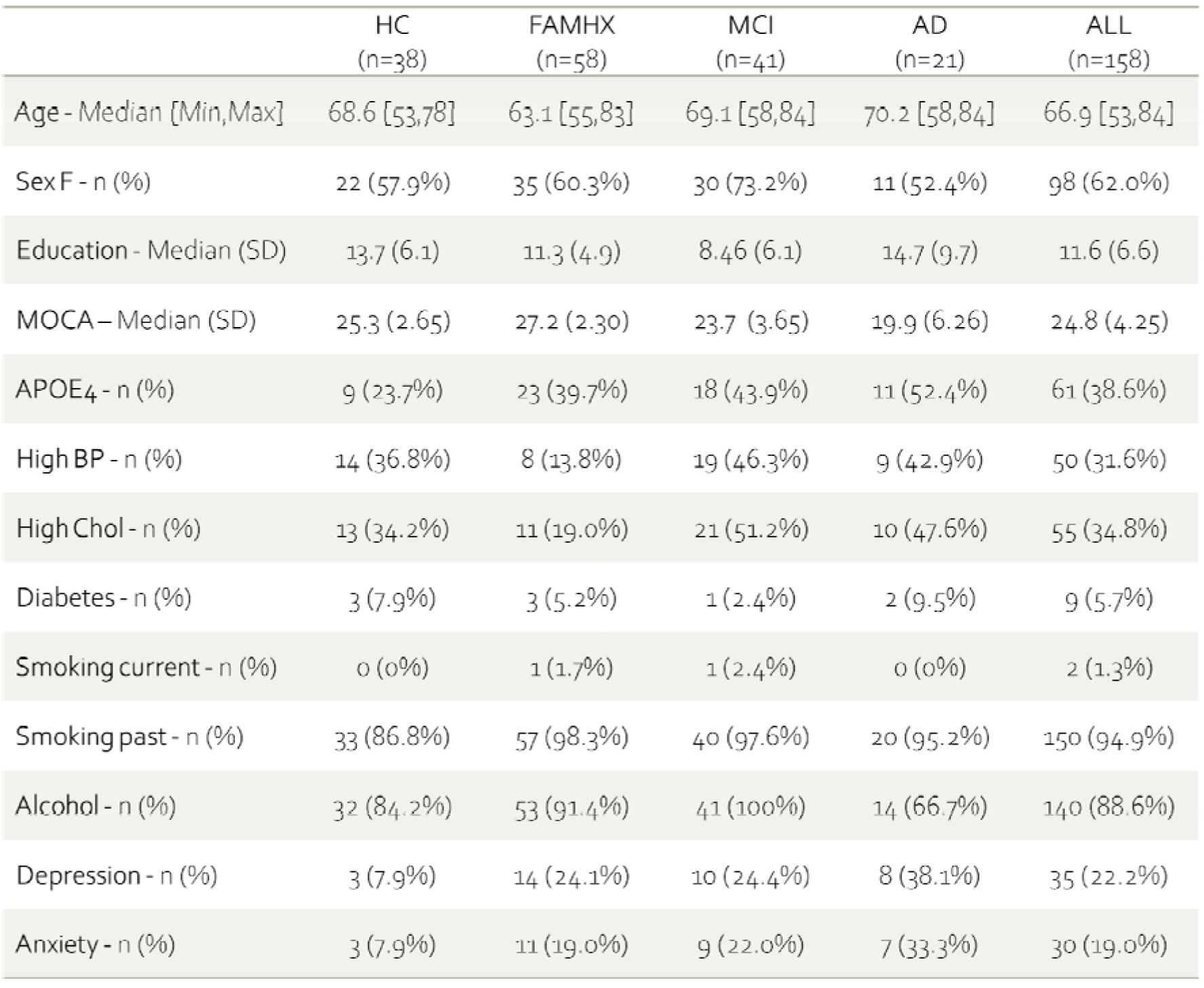
Demographic information of the 158 participants who passed QC and included in the study.

### 2.2. Demographic variables

Trained research assistants used questionnaires to gather information in four broad categories: cognitive, psychological, medical, and lifestyle. All this information will be collectively referred to as ‘demographic variables’ for the remainder of this study for the sake of readability.

- Cognition was assessed using the Montreal cognitive assessment (MOCA)^31^ and the repeatable battery for the assessment of neuropsychological status (RBANS)^32^, (see **supplementary methods** for additional information about the tests).
- For psychiatric variables, binary variables for depression and anxiety were used. A value of 1 indicates that an individual reported ever having an anxiety or depressive episode that required the care of a physician and pharmacological treatment, while a value of 0 indicates that they never had such an episode.
- We included information on medical history known to be comorbid with risk for AD or maladaptive aging, including hearing problems^33^, concussion^34^, transient ischemic attack^35^, brain injury^36^, headaches^37^, seizures^38^, heart disease^39^, liver disease^40^, kidney disease^41^, thyroid disease^42^, cancer^43^, arthritis^44^, neck/back problems^45^, and allergies^46^.
- We also examined modifiable lifestyle-related risk factors known to increase risk for AD; these included: alcohol use^47^, smoking^48^, drug use^49^, high blood pressure^50^, high cholesterol^51^, and diabetes^52^.

Because some variables had a limited amount (≤ 10.75%) of missing data (see **supplementary table 1**, we imputed missing values with Random Forest imputation using the missForest package (version 1.4) on R (version 3.5.1) for the 158 included individuals.

### 2.3. Imaging acquisition

Participants of the two datasets were scanned on the same 3T Siemens Tim Trio MRI scanner at the Douglas Research Center using a 32-channel head coil.

- We acquired whole brain magnetization prepared 2 rapid acquisitions by gradient echo (MP2RAGE) sequence at 1 mm isotropic resolution^53^. T1w uniform image (UNI; unbiased from B1 inhomogeneity, T2* and PD) was used for morphological characterization, and a map of the longitudinal spin-lattice relaxation time T1 was used for microstructural characterization of the myelin content. The MP2RAGE acquisition parameters are: inversion time (TI)1/TI2 = 700/2500 ms, echo time (TE)/repetition time (TR) = 2.91/5000 ms, flip angle (□)1/□2 = 4/5 deg, field-of-view (FOV) =256 x 256 mm2, 176 slices, 1 mm isotropic voxel dimensions. The image acquisition is accelerated in the phase encode direction by a GeneRalized Autocalibrating Partial Parallel Acquisition (GRAPPA) factor of 3, for a total scan time of 8 min 22 sec. QC of the T1w raw images were performed using a standardized procedure^54^ (https://github.com/CoBrALab/documentation/wiki/Motion-Quality-Control-Manual) to limit the well-known confounds of motion on downstream MRI-derived measurements^55,56^.
- A 3D multi-echo T2*-weighted gradient echo scan was acquired with the following parameters: 12 TE = [2.84, 6.2, 9.56, 12.92, 16.28, 19.64, 23, 26.36, 29.72, 33.08, 36.44, 39.8] ms, TR = 44 ms, bandwidth = 500 Hz/Px, □ = 15 deg, FOV = 180 x 192, 144 slices, 1 mm isotropic resolution, phase partial Fourier 6/8, GRAPPA acceleration factor = 2, for a total scan time of 9 min 44 sec. Estimation of the transverse T2* relaxation time is explained in **Section 2.6** and is considered a marker of iron content^57^.

While it is true that the correlation between T1 and T2* times with myelin and iron, respectively, are frequently acknowledged, the biological specificity to these mechanisms is far more complex. We delve into a more comprehensive description of the putative biological specificity underlying the observed signal in Discussion 4.3.

### 2.4. Image processing

#### 2.4.1. Preprocessing

The minc-bpipe-library pipeline (https://github.com/CobraLab/minc-bpipe-library) was employed to preprocess T1w images and performed N4 bias field correction^58^, standardized the FOV and head orientation, extracted the brain and created brain masks using BEaST^59^. QC at every step of the pipeline was performed (https://github.com/CoBrALab/documentation/wiki/Motion-Quality-Control-(QC)-Manual).

#### 2.4.2. T2* map extraction

Each GRE echo was denoised using an adaptive non-local means algorithm^24^ to improve the signal-to-noise ratio of the scans (see **Supplementary Figure 1**). The T2* exponential decay constant was estimated voxel-wise using the Levenberg-Marquardt *curve fit* function (https://github.com/CoBrALab/minc-toolkit-extras/blob/master/t2star_fit_simpleitk.py) using a single exponential model of the T2* decay equation:

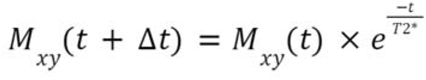

Where *M_xy_* is the transverse magnetization at time t.

QC of the T2* maps was performed to examine the scan quality based on motion artifacts and to remove scans with signal drop off (mostly in the temporal lobe) or spurious signal intensity variation throughout the map (see examples of good and bad quality T2* maps in **Supplementary Figure 2**).

#### 2.4.3. Morphometric and qMRI cortical feature extraction

All T1w UNI images were processed with the CIVET pipeline (version 2.1.1)^60,61^ to estimate vertex-wise CT and SA. Vertex-wise T1 and T2* values were sampled across the different depths with increments of 12.5% from the pial surface (“0% surface”) to the gray matter (GM) and white matter (WM) interface (“100% surface”) following similar approaches described by others^62,63^ and averaged. To correct for the T2* dependence on myelinated fiber orientation relative to , T2* values were residualized from the angle □ between the vertex normal and ^64^. All cortical measurements (morphometric and qMRI) were performed at each vertex of the 40,962 vertices per hemisphere and were spatially smoothed with a 30 mm surface-based heat diffusion kernel^65^. The medial wall was masked and therefore only 38,561 vertices per hemisphere were considered in the analyses. More details about the processing are available in **Supplementary methods**.

#### 2.4.4. Morphometric and qMRI hippocampal feature extraction

To derive voxel-wise volumetric measurements of the hippocampus, we used a python pipeline for deformation based morphometry (DBM), developed by the CoBrA Lab https://github.com/CobraLab/twolevel_ants_dbm. This technique uses the ANTs tools to obtain a minimally biased template using a group-wise registration strategy^66^, reproducing the approach employed in previous neurodegeneration studies conducted within our research group ^67^. We calculated voxel-wise volume differences by estimating the Jacobian determinant at each voxel from the individual non-linear displacement fields^68^. In this study, we used the relative Jacobian (J) determinant to account for overall differences in brain size (explicitly modeling the non-linear part of the deformations after linear scaling to the template). J values were log-transformed to aid statistical analysis: positive values indicate volumetric expansion whereas negative values indicated relative decrease volume.

To ensure that DBM outputs, T2* and T1 maps have voxel-wise correspondence across participants, we applied the transforms to T2* and T1 maps. A hippocampal mask was created on the average T1w brain to extract voxel-wise hippocampal information: J, T1 and T2* values (more information about the mask creation in **Supplementary methods** and **Supplementary figure 3**).

### 2.5. Statistics

#### 2.5.1. Non-negative matrix factorization

To integrate the multiple measures from morphometry and microstructure, we sought to use a multivariate method that detects covariance patterns that jointly describe all metrics used. To do this we leveraged non-negative matrix factorization (NMF) ^69^ as previously done in other publications in our group ^18,29,69–72^. NMF detects underlying patterns of covariation from complex data while promoting sparsity in the solution, resulting in spatially non-overlapping regions of covarying metrics.

This method decomposes an input matrix of vertices by subjects into two matrices: (i) representing spatial parcellation (vertices by components, **Figure 2A.1** and **Figure 2B.1**), and (ii) reflecting metrics for each subject within each component (components by subjects, **Figure 2A.2-3** and **Figure 2B.2-3**). We determined the number of components by assessing reconstruction stability and accuracy across different granularities as described in **Supplementary methods,** represented in **Supplementary Figures 4** and **5**, and in accordance with our previous work ^18,70,72^.

**Figure 2:**
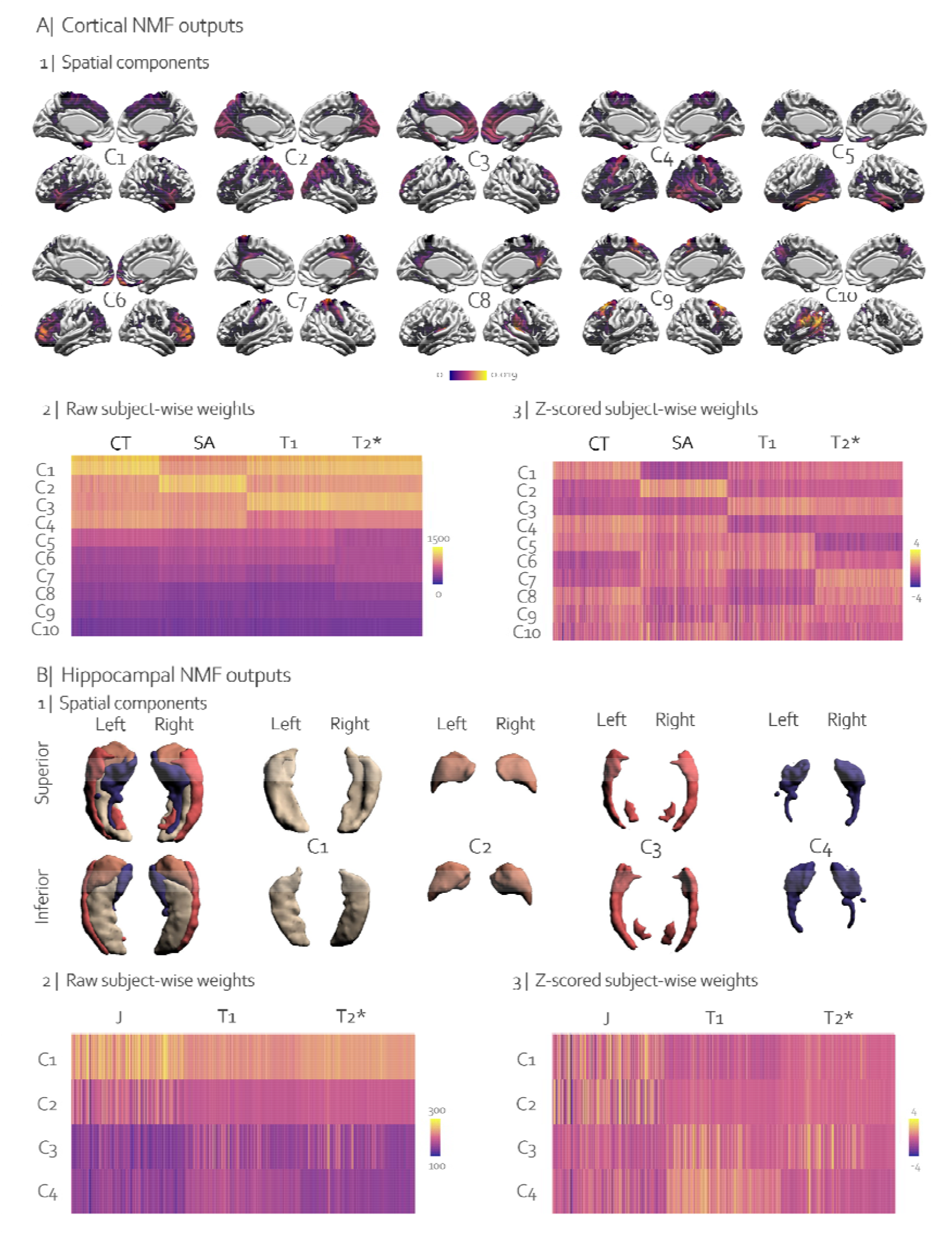
Visualization of the spatial components. **A | 1** Visualization of the 10 spatial components where each component is specific to a brain region. Component 1 is in the dorsomedial and superior temporal regions, component 2 in the occipital lobe, component 3 in the superior frontal cortex, component 4 in the auditory/motor cortices, component 5 in the inferior/medial temporal lobe, component 6 in the frontal lobe, component 7 in the cingulate/somatosensory regions, component 8 in the precuneus, component 9 is the premotor cortex and component 10 in the temporo-parietal junction. **A | 2** Raw subject-metric weights matrix, where each line corresponds to a spatial component, and each column represents a subject-metric weight. **A | 3** Standardized subject-metric matrix by z-scoring across rows (i.e per component) to better visualize which metric contributes the most to each component. For example, across our individuals, CT contributes the most to component 1 while SA contributes the most to component 2. **B | 1** Visualization of the 4 hippocampal components, where each component is specific to a hippocampal region. Component 1 is in the body and tail, component 2 in the head, component 3 in the lateral regions, and component 4 in the medial regions of the hippocampus. **B | 2** Raw subject-metric weights matrix and **B | 3** standardized subject-metric matrix by z-scoring across rows.

#### 2.5.2. Univariate analyses

To examine the relationship between NMF weights and our groups, we used ANCOVAs while correcting for age. Each component and metric were tested independently, totaling 52 models (4 * 10 cortical components + 3 * 4 hippocampal components). Subsequently, post-hoc Tukey Honestly Significant Difference (HSD) tests were employed to assess pairwise group differences. The resulting p-values from all models were corrected for False Discovery Rate (FDR) correction (q=0.05). Additionally, ANCOVA 95% confidence intervals for overall group differences are shown in **Figure 3**.

**Figure 3:**
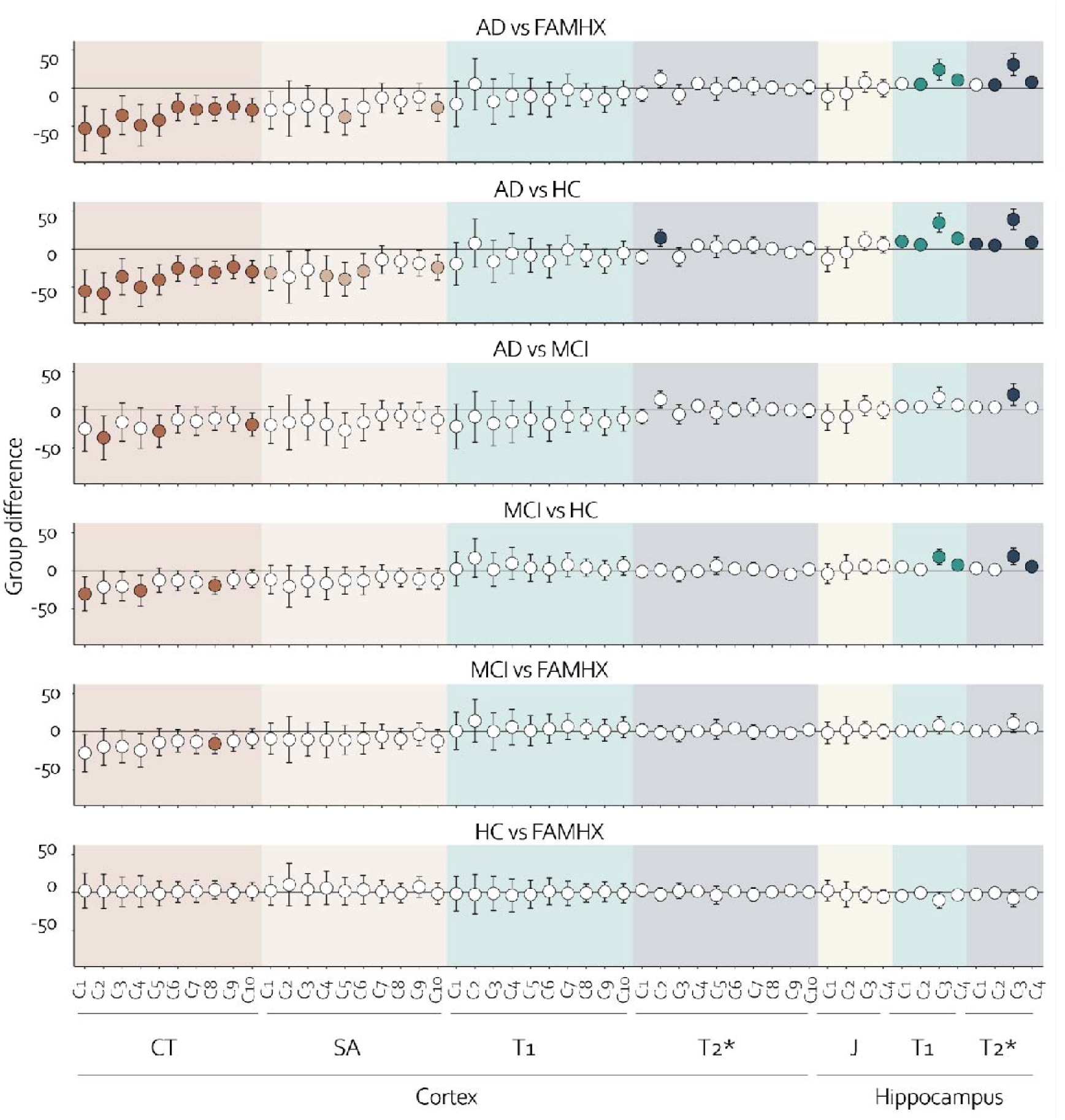
Pairwise NMF weights comparisons. NMF metric-wise and component-wise weights were compared pairwise across all combinations of groups The metrics included cortical CT, SA, T1, T2*, and hippocampal J, T1, and T2*. The mean group difference was indicated by circles, with gray circles representing non-significant differences, and colored circles representing significant differences after FDR correction across all p-values. The plots are color coded by metric and the 95% confidence interval of the group difference is represented by vertical bars.

#### 2.5.3. Investigating joint-brain and demographic signatures using partial least squares

Upon extracting our joint patterns of covariance in the MRI data, we examined their relevance to AD-related disease progression. To accomplish this we used partial least squares (PLS) analysis to relate the NMF-component individual metric weights to demographic signatures, as done in previous studies by our group^18,70–72^. PLS is a multivariate statistical technique commonly used to investigate the relationships between brain measures and demographic measures. This technique identifies patterns of covariance between two matrices: one matrix represents brain measures, and the other matrix includes demographic information. By decomposing these matrices into a set of orthogonal latent variables (LV), PLS analysis identifies the most prominent patterns of covariance between the two matrices. Each LV in PLS analysis represents a linear combination of the brain and demographic matrices providing a representation of brain regions and demographic measures that are most closely related. Here, our brain matrix included NMF weights, and the other matrix included demographic variables described in **Section 2.2**, and no information about the clinical grouping APOE4 status, age, sex and years of education of the individuals was provided. The goal of this approach was to capture a pattern of brain and demographic information which could explain the most covariance without giving information about our participant’s status. Our brain data included the NMF weights from the cortex (10 components x 4 metrics) and the hippocampus (4 components x 3 metrics) combined (i.e., 52 metrics for 158 participants). The values in the demographic matrices were z-scored prior to performing PLS (i.e., z-score the values of each demographic variable across all subjects). Each LV was tested statistically using permutation testing and split half analyses and variable contribution was assessed with bootstrap resampling following a similar protocol as in previous studies^18,28,67,73–76^.

Brain and demographic scores obtained from PLS analyses were further analyzed to determine if they were associated with key demographic and clinical variables not included in the PLS analysis. The relationship between PLS scores and demographic/clinical variables (group, age, sex, APOE4 and education) were assessed using ANCOVAs. Each model included all of these demographic/clinical variables simultaneously, resulting in a total of six models (one for each brain and demographic scores of the 3 LVs). Post-hoc Tukey Honestly Significant Difference (HSD) tests were performed to examine pairwise group differences. False discovery rate (FDR) corrected p-values can be found in **Supplementary table 2**. The goal of these analyses was to determine if PLS successfully captured, using a data-driven approach, patterns of brain and demographic that are clinically relevant in the context of AD.

### 2.6. Data and code availability statement

Raw data from this cohort can be obtained through collaborative agreement and reasonable request but is not publicly available due to the lack of informed consent by these human participants.

The code used to run NMF is available at https://github.com/CoBrALab/documentation/wiki/opNMF-for-vertex-data. For executing PLS, please refer to the instructions provided in https://github.com/CoBrALab/documentation/wiki/Running-PLS-in-MATLAB-with-cross-sectional-data. The repository containing all NMF and PLS outputs, as well as the main R scripts used to plot the figures in this paper, can be found at https://github.com/AurelieBussy/Cortex_hippocampus_Alzheimer_project.

## 3. Results

### 3.1. Demographics

The final sample after quality control included 158 participants, with a mean age of 66.9 years, an age range between 53 and 84 years and 62% of females (**Table 1**). The FAMHX group was significantly younger than the other groups (p < 0.001) and education was lower for the MCI group compared to HC and AD (p < 0.01).

### 3.2. NMF decomposition

#### 3.2.1. Cortical decomposition

The outputs of the cortical NMF analyses are shown in **Figure 2A**. Although we initially considered including 4 components because of its high stability score, we found that the corresponding spatial outputs were confined to the primary lobar regions (see **Supplementary Figure 6**). Because this did not offer an expansive representation of the cortex, we opted to investigate more components. Since NMF maintained a reasonable stability and accuracy for an increasing number of components, we proceeded with 10 components. Each component qualitatively demonstrated bilateral patterns in distinct cortical regions described in **Figure 2A**.

#### 3.2.2. Hippocampal decomposition

The outputs of the hippocampal stability analyses are shown in **Figure 2B**. We selected 4 components as per previous work examining hippocampal neuroanatomy^18^ and replicated here with our stability analysis. The 4 components of the hippocampus consisted of the body and tail (component 1), the head (component 2), the lateral areas (component 3), and the medial areas (component 4).

### 3.3. NMF components

#### 3.3.1. Cortical components

Each component can be described by their spatial pattern and their subject-wise and MRI metric-wise weights. For example, component 1 corresponds to the dorsomedial and superior temporal regions, high CT weights, low SA weights, and medium T1 and T2* weights. These values are apparent in both the raw subject-weight (**Figure 2 A| 2**) and standardized subject-weight matrices (**Figure 2 A| 3**). Accordingly, these patterns suggest that CT plays a greater contribution than SA in the spatial component 1.

However, comparing the contribution of a metric across different components (as shown in **Figure 2)** is not possible because the subject-wise weights are standardized per component. Nevertheless, we can compare the NMF weights of individuals within the same component and interpret the weights as raw values (e.g., average CT in each spatial region). To compare each metric across components, refer to **Supplementary Section 6** and **Supplementary figure 7.**

#### 3.3.2. Hippocampal components

In the same way, each hippocampal component can be described with the subject-metric weights. J mostly contributed in component 1 and 2 patterns (hippocampal body and head) while T1 contributed the most to component 4 (medial hippocampus) and both T1 and T2* contributed more than J in component 3 (lateral hippocampus).

### 3.4. Group differences

We assessed whether our NMF components could differentiate our groups by using cortical and hippocampal weights, as shown in **Figure 3**. Comparing AD to FAMHX, we observed lower CT throughout the brain, reduced SA in component 5 and 10 (which correspond to the temporal lobe and the temporo-parietal junction), and longer T1 and T2* in components 2, 3, and 4 of the hippocampi (which correspond to the head, lateral, and medial regions). A similar but more widespread pattern was observed when comparing AD and HC, with reductions in SA observed in more regions of the cortex and longer in T1 and T2* seen throughout the entire hippocampus. Additionally, T2* was longer in the occipital regions of individuals with AD vs HC. Comparing AD and MCI, we observed fewer significant differences, but we still found lower CT in the occipital, temporal, and temporo-parietal junction, as well as longer T2* in the lateral region of the hippocampus. Comparing MCI and HC, we found lower CT in the dorsomedial and superior temporal regions, in the auditory/motor cortices, and in the precuneus, as well as longer T1 and T2* in the lateral and medial regions of the hippocampus. When comparing MCI and FAMHX, we only observed lower CT in the precuneus, and no significant differences were found between FAMHX and HC. Our results suggest that cortical morphometry and hippocampal microstructure are the metrics the most sensitive to AD progression, making it possible to distinguish between MCI and controls for example. Indeed, individuals with AD showed greater cortical thickness and surface reduction as well as longer hippocampal T1 and T2* compared to those with MCI, who in turn demonstrated significant differences from the control group.

### 3.5. Brain and demographic relationships

#### 3.5.1. Latent variables

Our PLS results demonstrated three significant LVs, explaining 76.9% (p<0.0001), 7.5% (p<0.001) and 5.2% (p<0.001) of the covariance respectively. The results of the first LV are shown in **Figure 4** (LV2 and LV3 in **Supplementary Figure 8 and 10**; both LVs predominantly captured patterns associated with sex but showed no significant correlation with age or AD progression). **Figure 4A** shows the brain pattern, describing a lower CT, SA in the entire brain, shorter T2* in the dorsomedial and superior temporal regions, superior frontal cortex and in the premotor cortex and longer T2* in the occipital lobe. We found a decreased J in the body and tail, an increased J in the most lateral region as well as longer T1 and T2* values in all regions of the hippocampus. Importantly, we observed that the hippocampal microstructure exhibits stronger contributions to AD-relevant variables compared to its volumes. We observed that cortical morphometry and hippocampal microstructure were linked to a pattern of past smoking consumption, high blood pressure (BP) and cholesterol, lower scores in all cognitive domains, and higher anxiety levels (**Figure 4B**).

**Figure 4:**
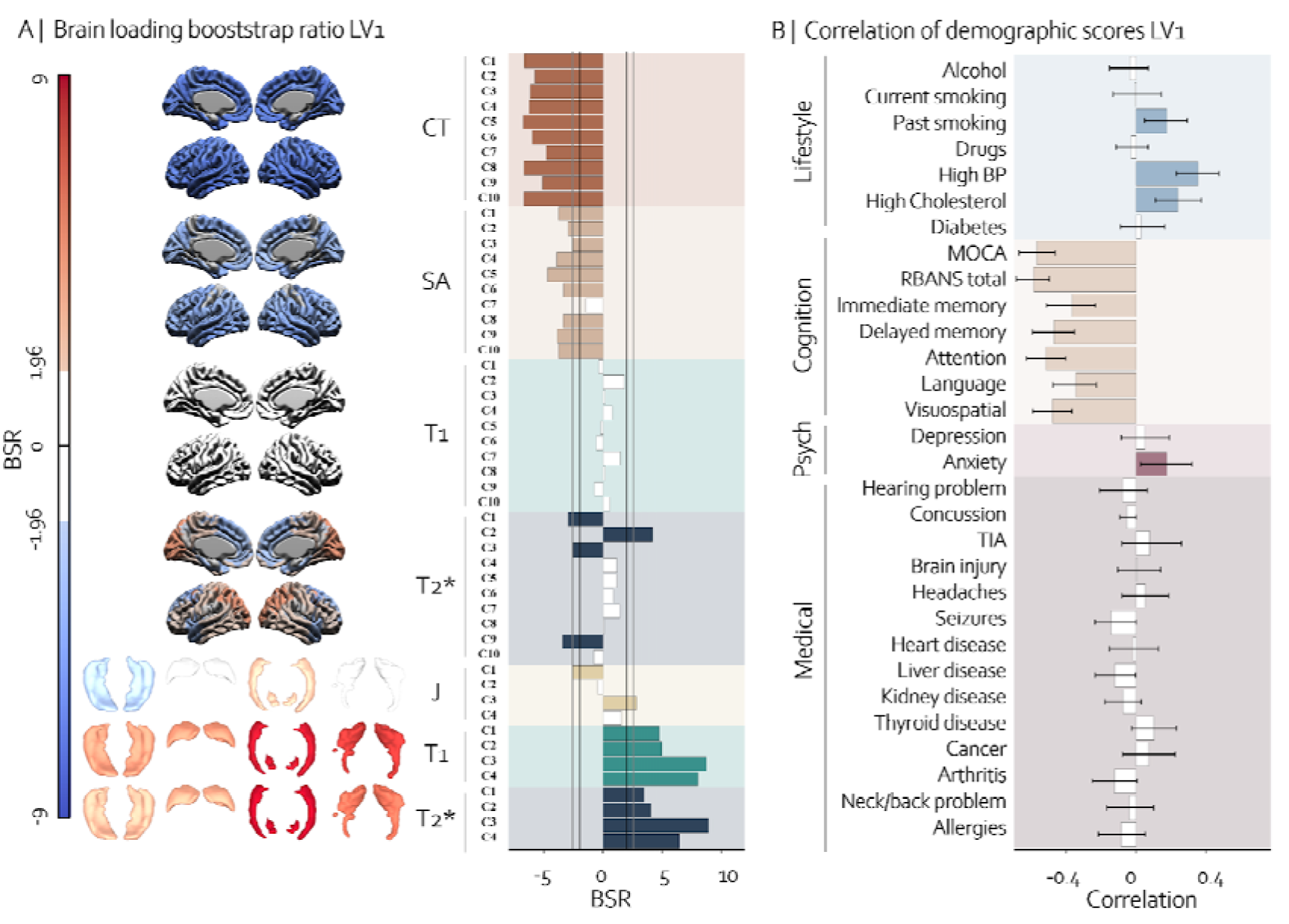
Relationship between brain and cognitive, psychological, medical and lifestyle information of the LV1. The first LV explained 76.9% of the covariance between brain and cognitive, psychological, medical, and lifestyle information. The brain pattern on the left **(A)** illustrates spatially the contribution of each metric to the pattern with bootstrap ratio (BSR) values on each brain structure. Blue color indicates negative BSR values and red indicates positive BSR values. Components with absolute BSR values higher than 1.96 are colored to show significant contribution. The bar plot in the middle of the figure shows more precisely the BSR values for each component, with black vertical lines representing a BSR of 1.96 (equivalent to p=0.05) and gray lines representing a BSR of 2.58 (equivalent to p=0.01). On the right-hand side **(B)**, we show the cognitive, psychological, medical, and lifestyle patterns associated with the brain pattern on the left. Bars are colored if they are significant (when error bars do not cross zero) and are white if non-significant. The brain pattern is associated with past smoking consumption, high BP and cholesterol, lower cognitive scores, and higher anxiety.

**Figure 5:**
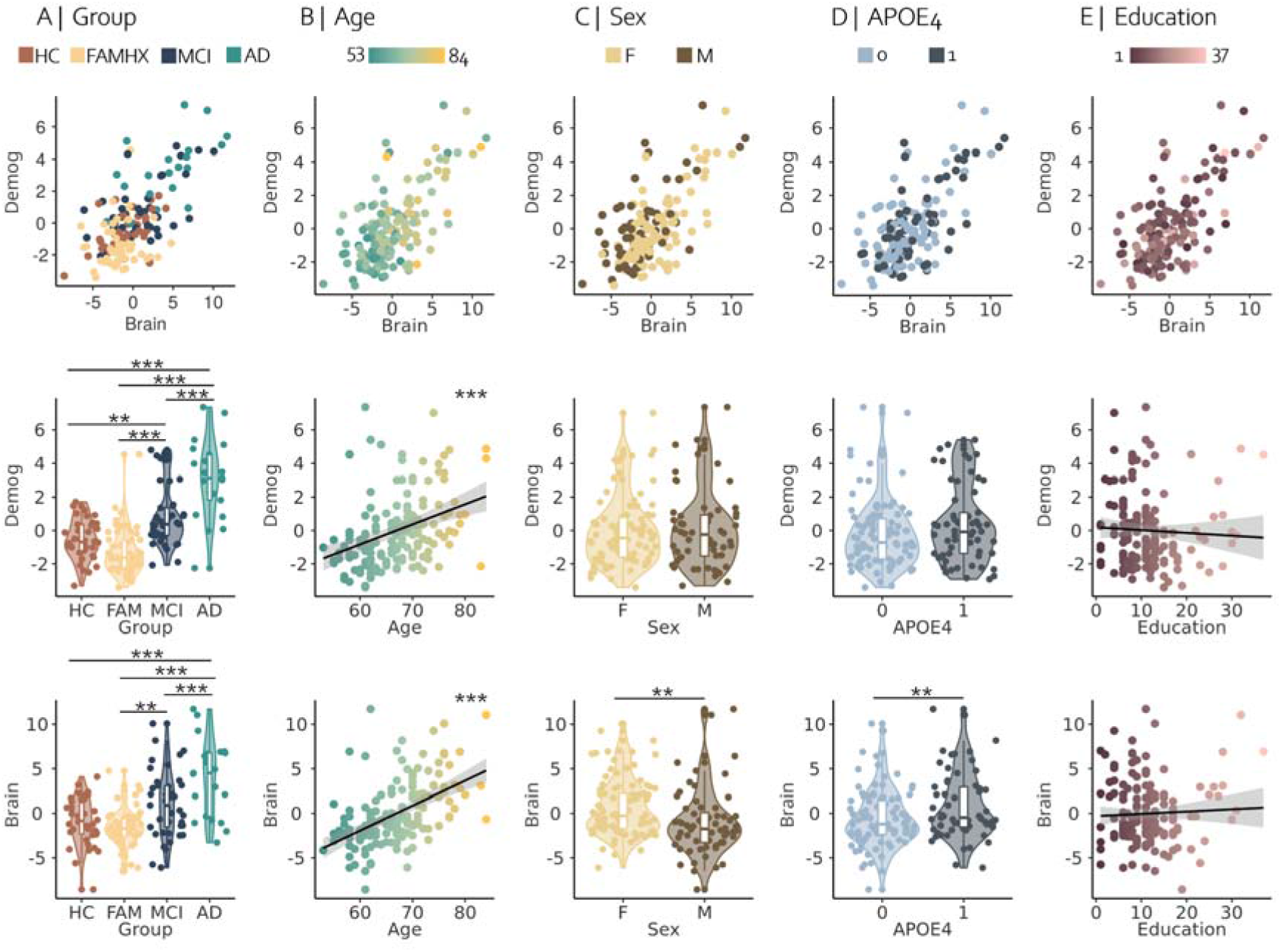
Post-hoc analyses of the PLS brain and demographic scores of LV1. **(A)** pairwise group comparisons of the brain and demographic scores. Results of the linear models illustrating the brain and demographic relationships with age **(B)**, sex **(C)**, APOE4 status **(D)** and education **(E)**. **\*** p < 0.05, **\*\*** p < 0.01, **\*\*\*** p < 0.001 after FDR correction across all p-values.

#### 3.5.2. Brain/demographic pattern sensitive of disease progression, age and genetic risk

Lastly, we explored how age, group, APOE4 status, sex, and education related to the brain and demographic patterns of the LV1. Similar analyses were performed for the LV2 and LV3 pattern (**Supplementary Figure 9 and 11**). We observed significant group differences between each pair of groups, except for HC vs FAMHX in the brain and HC vs FAMHX and HC vs MCI in the demographic scores. Our findings indicated that both brain and demographic were positively associated with age while only brain scores were significantly associated with female sex. Additionally, APOE4 carriers demonstrated a stronger expression of the brain pattern than non-carriers. No significant effect of education was observed in relation to brain and demographic scores. In summary, the results demonstrate a significant association between the highlighted brain and demographic patterns in LV1 and AD progression, particularly among older individuals, females, and carriers of the APOE4 allele.

## 4. Discussion

In this study, we examined if brain morphometry and qMRI markers of iron and myelin could be sensitive to AD progression and AD-related risk factors. First, we took a data-driven approach to parcellate the cortex and the hippocampus using NMF by integrating multimodal MRI data to take advantage of the complementary information conveyed by these indices. Overall, our results suggest that combining multi-contrast MRI metrics is critical for gaining a more nuanced understanding of the properties of the brain and is well-adapted to identify their differential susceptibility to adaptive and maladaptive aging and AD.

### 4.1. Brain patterns associated with AD

Our analysis of pairwise group comparisons of NMF weighted demonstrated that cortical thinning was apparent throughout the brain when we compared AD vs FAMHX or AD vs HC. However, the cortical signature of AD is often characterized by regionally specific cortical thinning related to symptom severity in the temporal and frontal regions^77,78^, while our results did not necessarily show stronger effects in these regions. A previous study from our group using similar methodologies in a healthy aging population demonstrated a widespread association between cortical SA and performance across cognitive domains in midlife ^71^. Although not as widespread, our results demonstrated that some level of SA reduction in the temporal/parietal lobe was observable in AD relative to HC^79^. Further, our results demonstrated that CT has larger effect size than SA in the LV. Notably, significant differences were observed between MCI participants and controls for CT, although SA showed significance solely in the more advanced disease stage comparisons (AD vs. controls). These findings underscore the greater sensitivity of CT to AD progression when compared to SA^80^. Overall, even though FAMHX carry a higher genetic risk, their morphological patterns seemed to be similar with those from the HC group. The resemblance between these groups may be attributable to the lower age of the FAMHX relative to controls.^30^.

Surprisingly, contrary to our cortical morphometry findings, no hippocampal volume differences were found between groups in our univariate analyses. However, a volume reduction of the hippocampal body and tail, and an increase in volume in the hippocampal lateral region were found in LV1. These results suggest that while hippocampal volume did not show a specific relationship to a univariate group effect, it may be more useful to model this variation using methods that capture disease spectrum rather than simply searching for group differences. This is supported by the specificity of our demographic patterns (discussed in **Section 4.2**). Interestingly, the hippocampal parcellation obtained in this study was similar to the one we previously reported^18^, suggesting that this organization is a consistent microstructural pattern in the human hippocampus.

Cortical T1 and T2* did not differ between our groups, while hippocampal T1 and T2* did. Therefore, we hypothesize that the cortical thinning might not be driven by intra-cortical demyelination or iron accumulation but rather by neuronal death^81^. This further suggests that lower myelin content associated with AD may initiate in the hippocampus prior to manifesting in the cortex. This is in line with previous studies showing a specific myelin decrease in the hippocampus using magnetization transfer measurements in MCI individuals^82^ and in AD individuals compared to controls^9^. Using T1w/T2w ratio to estimate intracortical myelin, another study demonstrated that hippocampal demyelination was consistently associated with AD progression^83^. Previous research showed lower T2* in the hippocampus of those with AD, indicating an increase in iron levels^84^. However, our study revealed the opposite pattern, with disease progression associated with a longer hippocampal T2*. Further discussion on these findings can be found in **Section 4.3**.

### 4.2. Demographic alterations associated with AD

Our results demonstrated demographic variables that are well-described as risk factors for AD; namely: lower cognitive scores, higher anxiety, BP, cholesterol and smoking^85^, were associated with our brain alterations and overall AD progression.

About 70% of AD patients present anxiety symptoms^86^ which have been linked to a higher risk ratio of converting to MCI or dementia^87^, worse mini mental state examination (MMSE) scores and younger age at onset^88,89^. Interestingly, in a healthy aging population, anxiety disorders have been previously related to a longer of T1 values in the hippocampus^90^. This is consistent with our results showing a longer hippocampal T1 associated with higher anxiety and AD progression. We also found that the pattern of increased anxiety was significantly related to cortical thinning across the cortex. Similarly, previous research reported a relationship between higher levels of anxiety symptoms and reduced thickness in several cortical regions^91^.

A negative correlation between arterial hypertension and myelin content has been observed in the WM^90^. Late-life cortical and WM atrophy have been found to be linked with hypertension during early adulthood^92^. In line with these findings, our study showed a similar effect, where higher BP was associated with myelin reduction and cortical thinning^93^. Furthermore, our findings indicated a significant association between the brain pattern, disease progression and hypertension, aligning with existing research in this field^94,95^.

Elevated blood cholesterol levels have been reported to increase Aβ production in the brain^96–98^. Conversely, drugs that reduce blood cholesterol have been shown to lower the risks of developing AD^99,100^. It’s worth noting that the blood-brain barrier (BBB) typically prevents any exchange between the brain and the cholesterol, which means that most brain cholesterol comes from local synthesis. Further, it has been shown that the hippocampus is a brain region particularly susceptible to BBB breakdown^101^, and as a result, there may be early alterations in the level of hippocampal cholesterol in AD progression. However, while most studies report brain cholesterol increases related to aging and AD^98,102,103^, others found the opposite effect^104^.

The identification of the APOE4 allele as a crucial genetic risk factor for AD is in accordance with the involvement of cholesterol in AD’s pathogenesis^51,104,105^. Notably, in APOE4 carriers, ApoE exhibits reduced binding capacity and transport affinity for lipids^106,107^, which may decrease the transfer of cholesterol from astrocytes to neurons, eventually leading to neuronal apoptosis. This aligns with our findings, which demonstrated that APOE4 carriers were more heavily loaded in the pattern of brain and lifestyle risk factor association that we observed. While there is a need to clarify the exact relationship between brain cholesterol levels and AD, altered hippocampal cholesterol could explain the hippocampal demyelination pattern related to being APOE4 carriers.

Finally, research has shown that smoking was associated with an increased risk of dementia and AD^108,109^. Smoking has also been linked to reduced brain volume and atrophy in specific cortical regions, including the frontal, occipital, and temporal lobes^110,111^. Even when taking into account the amount of tobacco smoked over their lifetime, individuals who currently smoke had greater hippocampal atrophy than those who never smoked or had quit smoking in the past^112^. Smokers have also been found to have a greater rate of atrophy in regions that are affected in the early stages of AD compared to non-smokers^113^. Furthermore, cigarette smoke is known to trigger the production of endogenous oxidants by activating the immune response pathway associated with inflammation^114,115^. Significant positive relationships between R2* and smoking were found in certain brain regions such as the basal ganglia, but not in the hippocampus^90^. Here, we found a longer hippocampal T2* being associated with past smoking consumption. This difference of findings could be explained by the fact that our pattern was found in the context of AD progression while Trofimova et al.^90^ included healthy participants. Although R2* is commonly interpreted as being solely related to iron content, in **Section 4.3**, we discuss how other mechanisms linked to AD pathology can influence these metrics.

### 4.3. Specificity of qMRI metrics

We employed two qMRI methods giving important tissue relaxation times, namely T1 and T2* maps. T1 maps offer insights into the longitudinal relaxation time constant (T1) at each voxel, influenced by factors like myelin^116,117^, iron^118^, and proton density (PD)^119^. Increased myelin and iron content reduce T1, while increased water content extends T1, particularly observed in subcortical gray matter where T1 variation is linked to myelin content^120^. Complementing T1 mapping, T2* relaxation time reflects dephasing due to molecular interactions and local magnetic field inhomogeneities^121^, primarily influenced by iron content^57,122,123^.

Few validation studies have been performed in the context of diseased tissue. For example, T1 has been almost only validated compared to myelin in multiple sclerosis brains^124–127^ or in animal models with experimentally induced demyelination^128,129^. T2* (or R2*) has been validated against iron in a larger range of applications, from cardiac studies^130^, hepatic studies^131^, to neurological studies in both multiple sclerosis^132,133^ and in healthy individuals^123,134^. However, it is currently unclear what T1 and T2* metrics reflect in an AD brain where other microstructural changes, such as neuronal loss and increased amyloid and tau accumulation, occur simultaneously.

Interpretation of T2* is complex as it combines the effects of transverse relaxation T2 and magnetic susceptibility T2’, where increased water content increases T2 and increased iron content decreases 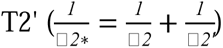. While several studies have reported an extended T2 in the hippocampus of individuals with AD^135–137^, others have observed the opposite effect^138^ or no change at all^139^. Longer hippocampal T2 in the AD group has been postulated to be linked to increased water reflecting tissue damage^16^. Indeed, in neurodegenerative disorders, the increased water content in degenerating tissue can also affect MRI relaxation times and reduce R2, which opposes the effect of iron^135^. Therefore, in AD, the increased water content may make it difficult to detect increased iron levels with T2*^140^.

Our results demonstrated significantly longer T1 and T2* in the hippocampus of individuals with AD compared to HC. T1 is primarily linked to myelin content, where a longer in T1 generally indicates lower myelin content^116^. A reduction of glial cell density, including astrocytes and oligodendrocytes could lead to a reduction of ferritin leading to a decreased iron^141^. Further, we know that T2* mostly reflects the amount of ferritin^142^ Therefore, we postulate that our findings principally captures a reduction in hippocampal tissue integrity, decreased myelin, increased water content and iron reduction.

### 4.4. Role of myelin in AD progression

Starting with the pioneering work of Bartzokis’ ^10,11^, myelin deterioration has been suggested as being an important factor in the progression of AD. Several arguments highlight the link between myelin and AD. First, humans are the only animal susceptible to AD pathology. Indeed, even if some nonhuman primates and dogs develop amyloid, they do not present tau or dementia-like symptoms^143^.

Supporting the fact that amyloid might not be the main molecule triggering cognitive decline, combined data from multiple trials has demonstrated that reducing amyloid levels do not significantly enhance cognitive function^144^. Another important criticism is that both neuropathological and PET data reveal substantial evidence of Aβ pathology in older individuals who do not necessarily exhibit cognitive impairment^145^. Altogether, the presence of Aβ deposition without any cognitive impairment, along with the reduction of Aβ levels without any cognitive improvement, raises significant concerns regarding the validity of Aβ as a causal factor of the clinical symptoms of AD^146,147^.

Interestingly, there is a noticeable similarity between the pattern of neurofibrillary tangles (NFT) changes observed in AD and the reverse order of cortical myelination^148^. Indeed, certain brain regions, such as the prefrontal cortex and association areas such as the parietal and temporal lobes, which are characterized by late myelination, are particularly susceptible to the development of amyloid and NFT. Late myelinated regions which have thinner myelin sheaths^149^ are more susceptible to degeneration^150^. Conversely, heavily and early myelinated regions of the brain, such as the primary motor and sensory areas, appear to be more resistant to the disease^149^. In our results, while we did not find significant cortical T1 variation related to the disease progression, we found high sensitivity of hippocampal demyelination to disease progression. These findings are consistent with the hippocampus being one of the first regions impacted by AD pathology^151^.

Studies have shown that higher amyloid deposition assessed by PET is associated with lower T2* in the cortex^84^. Therefore, our findings indicating a lower T2* in the dorsomedial and superior temporal regions, superior frontal cortex, and premotor cortex are consistent with an increased pathological burden in those areas. Notably, these regions correspond to late myelinated regions, which aligns with the theory that late myelinated regions are at higher risk of developing AD pathology. In contrast, we observed an opposite pattern in early myelinated regions such as the occipital lobe, which demonstrated a longer T2* value associated with our brain pattern. The cause of this T2* lengthening is unclear, since it could be related to the tissue water content, a myelin reduction, death of glial cells, an iron decrease or potentially a methodological limitation.

### 4.5. Methodological limitations

The MP2RAGE sequence uses small flip angles and an adiabatic inversion pulse to significantly mitigate the impact of □*1*^+^on resulting T1 maps^53^. Despite this, residual biases linked to □*1*^+^ may persist, particularly near inferior temporal and frontal lobes. While proposed solutions involve acquiring additional □*1*^+^ maps to correct for these inhomogeneities to enhance quantitative T1 mapping^152^, our study lacked □*1*^+^ map acquisition, leaving the possibility that our results may be influenced by the presence of □*1*^+^ inhomogeneity residuals. However, since our results demonstrate good consistency between both hemispheres, we postulate that our results are not driven by this residual inhomogeneity.

Fiber orientation in relation to the main magnetic field □*_0_* affects T2* relaxation times in WM^153^. To reduce angle dependency in cortical T2*, we applied a suggested method^64^. However, estimating myelinated fiber orientation in the complex-shaped hippocampus proved impractical, and advanced metrics like diffusion tensor imaging were unavailable^154,155^. Due to the intricate nature of T2* correction, clinical papers often omit this step in applications. Consequently, we believe that our results are comparable to most findings in the field.

Unfortunately, our exploration of age and AD progression was constrained to cross-sectional data. We acknowledge the inherent limitations of this approach and recognize that future studies using longitudinal data would be necessary to validate our findings.

## Future research

This work has several potential avenues for future investigation. First, to further validate our interpretation, additional qMRI markers, such as PD and magnetization transfer metrics, could be used alongside T2* and T1. PD, in particular, could provide valuable information about the water content in the tissue. Second, the relationship between qMRI and AD pathology is not yet well characterized, and thus, including amyloid and tau PET scans alongside qMRI protocols would shed light on the impact of these pathological molecules on qMRI metrics. Finally, a longitudinal dataset with both PET and qMRI measurements in preclinical AD individuals would be ideal to determine the interplay between myelin, iron, amyloid, and tau.

## 5. Conclusion

In conclusion, by using qMRI to investigate the underlying biological processes that could drive morphological changes, our study suggests that hippocampal T1 and T2* could serve as potential biomarkers for AD. Further, significant associations between certain risk factors and AD-related brain pattern alterations demonstrate that public health initiatives aimed at reducing smoking, cholesterol levels, blood pressure, and anxiety in the population should be expanded to slow the progression of AD.

## Funding

**Dr Bussy** received support from the Alzheimer Society of Canada and Healthy Brains Healthy Lives. **Dr Chakravarty** is funded by the Weston Brain Institute, the Canadian Institutes of Health Research, the Natural Sciences and Engineering Research Council of Canada, and Healthy Brains Healthy Lives (a Canada First Research Excellence Fund project) and receives salary support from Fondation de Recherches Santé Québec. **Dr Patel** is funded through the Fonds de Recherche du Québec - Santé (Doctoral Training).

## Competing interests

The authors report no competing interests.

## Notes

### Competing Interest Statement

The authors have declared no competing interest.

## References

1. Rajan KB, Weuve J, Barnes LL, McAninch EA, Wilson RS, Evans DA. Population estimate of people with clinical Alzheimer’s disease and mild cognitive impairment in the United States (2020-2060). Alzheimers Dement. 2021;17(12):1966–1975.

2. Ito K, Ahadieh S, Corrigan B, et al. Disease progression meta-analysis model in Alzheimer’s disease. Alzheimers Dement. 2010;6(1):39–53.

3. van Dyck CH, Swanson CJ, Aisen P, et al. Lecanemab in Early Alzheimer’s Disease. N Engl J Med. 2023;388(1):9–21.

4. Tan CC, Yu JT, Wang HF, et al. Efficacy and safety of donepezil, galantamine, rivastigmine, and memantine for the treatment of Alzheimer’s disease: a systematic review and meta-analysis. J Alzheimers Dis. 2014;41(2):615–631.

5. Leng F, Hinz R, Gentleman S, et al. Neuroinflammation is independently associated with brain network dysfunction in Alzheimer’s disease. Mol Psychiatry. 2023;28(3):1303–1311.

6. Bateman RJ, Xiong C, Benzinger TLS, et al. Clinical and biomarker changes in dominantly inherited Alzheimer’s disease. N Engl J Med. 2012;367(9):795–804.

7. Granziera C, Wuerfel J, Barkhof F, et al. Quantitative magnetic resonance imaging towards clinical application in multiple sclerosis. Brain. 2021;144(5):1296–1311.

8. Eck BL, Yang M, Elias JJ, et al. Quantitative MRI for Evaluation of Musculoskeletal Disease: Cartilage and Muscle Composition, Joint Inflammation, and Biomechanics in Osteoarthritis. Invest Radiol. 2023;58(1):60–75.

9. Moallemian S, Salmon E, Bahri MA, et al. Multimodal imaging of microstructural cerebral alterations and loss of synaptic density in Alzheimer’s disease. Neurobiol Aging. 2023;132:24–35.

10. Bartzokis G, Lu PH, Mintz J. Human brain myelination and amyloid beta deposition in Alzheimer’s disease. Alzheimers Dement. 2007;3(2):122–125.

11. Bartzokis G. Alzheimer’s disease as homeostatic responses to age-related myelin breakdown. Neurobiol Aging. 2011;32(8):1341–1371.

12. Liu JL, Fan YG, Yang ZS, Wang ZY, Guo C. Iron and Alzheimer’s Disease: From Pathogenesis to Therapeutic Implications. Front Neurosci. 2018;12. doi:10.3389/fnins.2018.00632

13. Papuć E, Rejdak K. The role of myelin damage in Alzheimer’s disease pathology. Arch Med Sci. 2020;16(2):345–351.

14. Depp C, Sun T, Sasmita AO, et al. Myelin dysfunction drives amyloid-β deposition in models of Alzheimer’s disease. Nature. 2023;618(7964):349-357.

15. Bartzokis G, Sultzer D, Mintz J, et al. In vivo evaluation of brain iron in Alzheimer’s disease and normal subjects using MRI. Biol Psychiatry. 1994;35(7):480–487.

16. Raven EP, Lu PH, Tishler TA, Heydari P, Bartzokis G. Increased iron levels and decreased tissue integrity in hippocampus of Alzheimer’s disease detected in vivo with magnetic resonance imaging. J Alzheimers Dis. 2013;37(1):127–136.

17. Ward RJ, Zucca FA, Duyn JH, Crichton RR, Zecca L. The role of iron in brain ageing and neurodegenerative disorders. Lancet Neurol. 2014;13(10):1045–1060.

18. Patel R, Steele CJ, Chen AGX, et al. Investigating microstructural variation in the human hippocampus using non-negative matrix factorization. Neuroimage. 2020;207:116348.

19. Schuff N, Woerner N, Boreta L, et al. MRI of hippocampal volume loss in early Alzheimer’s disease in relation to ApoE genotype and biomarkers. Brain. 2009;132(Pt 4):1067–1077.

20. Jack CR Jr, Barkhof F, Bernstein MA, et al. Steps to standardization and validation of hippocampal volumetry as a biomarker in clinical trials and diagnostic criterion for Alzheimer’s disease. Alzheimers Dement. 2011;7(4):474–485.e4.

21. Jack CR Jr, Petersen RC, Xu Y, et al. Rate of medial temporal lobe atrophy in typical aging and Alzheimer’s disease. Neurology. 1998;51(4):993–999.

22. Duara R, Loewenstein DA, Potter E, et al. Medial temporal lobe atrophy on MRI scans and the diagnosis of Alzheimer disease. Neurology. 2008;71(24):1986–1992.

23. Visser PJ, Verhey FRJ, Hofman PAM, Scheltens P, Jolles J. Medial temporal lobe atrophy predicts Alzheimer’s disease in patients with minor cognitive impairment. J Neurol Neurosurg Psychiatry. 2002;72(4):491–497.

24. Manjón JV, Coupé P, Martí-Bonmatí L, Collins DL, Robles M. Adaptive non-local means denoising of MR images with spatially varying noise levels. J Magn Reson Imaging. 2010;31(1):192–203.

25. Rombouts SARB, Scheltens P, Kuijer JPA, Barkhof F. Whole brain analysis of T2* weighted baseline FMRI signal in dementia. Hum Brain Mapp. 2007;28(12):1313–1317.

26. Tullo S, Patel R, Devenyi GA, et al. MR-based age-related effects on the striatum, globus pallidus, and thalamus in healthy individuals across the adult lifespan. Hum Brain Mapp. 2019;40(18):5269–5288.

27. Bussy A, Plitman E, Patel R, et al. Hippocampal subfield volumes across the healthy lifespan and the effects of MR sequence on estimates. Neuroimage. 2021;233:117931.

28. Bussy A, Patel R, Plitman E, et al. Hippocampal shape across the healthy lifespan and its relationship with cognition. Neurobiol Aging. 2021;106:153–168.

29. Parent O, Bussy A, Devenyi GA, et al. Assessment of white matter hyperintensity severity using multimodal magnetic resonance imaging. Brain Commun. 2023;5(6):fcad279.

30. Tremblay-Mercier J, Madjar C, Das S, et al. Open science datasets from PREVENT-AD, a longitudinal cohort of pre-symptomatic Alzheimer’s disease. NeuroImage Clin. 2021;31(102733):102733.

31. Nasreddine ZS, Phillips NA, Bédirian V, et al. The Montreal Cognitive Assessment, MoCA: a brief screening tool for mild cognitive impairment. J Am Geriatr Soc. 2005;53(4):695–699.

32. Randolph C, Tierney MC, Mohr E, Chase TN. The Repeatable Battery for the Assessment of Neuropsychological Status (RBANS): Preliminary Clinical Validity. J Clin Exp Neuropsychol. 1998;20(3):310–319.

33. Uhlmann RF, Larson EB, Koepsell TD. Hearing impairment and cognitive decline in senile dementia of the Alzheimer’s type. J Am Geriatr Soc. 1986;34(3):207–210.

34. Guskiewicz KM, Marshall SW, Bailes J, et al. Association between recurrent concussion and late-life cognitive impairment in retired professional football players. Neurosurgery. 2005;57(4):719–726; discussion 719-726.

35. Liu W, Wong A, Au L, et al. Influence of Amyloid-β on Cognitive Decline After Stroke/Transient Ischemic Attack: Three-Year Longitudinal Study. Stroke. 2015;46(11):3074–3080.

36. Sivanandam TM, Thakur MK. Traumatic brain injury: a risk factor for Alzheimer’s disease. Neurosci Biobehav Rev. 2012;36(5):1376–1381.

37. Qu H, Yang S, Yao Z, Sun X, Chen H. Association of Headache Disorders and the Risk of Dementia: Meta-Analysis of Cohort Studies. Front Aging Neurosci. 2022;14:804341.

38. Amatniek JC, Hauser WA, DelCastillo-Castaneda C, et al. Incidence and predictors of seizures in patients with Alzheimer’s disease. Epilepsia. 2006;47(5):867–872.

39. de la Torre JC. How do heart disease and stroke become risk factors for Alzheimer’s disease? Neurol Res. 2006;28(6):637–644.

40. Estrada LD, Ahumada P, Cabrera D, Arab JP. Liver Dysfunction as a Novel Player in Alzheimer’s Progression: Looking Outside the Brain. Front Aging Neurosci. 2019;11:174.

41. Murray AM. Cognitive impairment in the aging dialysis and chronic kidney disease populations: an occult burden. Adv Chronic Kidney Dis. 2008;15(2):123–132.

42. van Osch LADM, Hogervorst E, Combrinck M, Smith AD. Low thyroid-stimulating hormone as an independent risk factor for Alzheimer disease. Neurology. 2004;62(11):1967–1971.

43. Musicco M, Adorni F, Di Santo S, et al. Inverse occurrence of cancer and Alzheimer disease: a population-based incidence study. Neurology. 2013;81(4):322–328.

44. McGeer PL, Schulzer M, McGeer EG. Arthritis and anti-inflammatory agents as possible protective factors for Alzheimer’s disease: a review of 17 epidemiologic studies. Neurology. 1996;47(2):425–432.

45. Wong AYL, Karppinen J, Samartzis D. Low back pain in older adults: risk factors, management options and future directions. Scoliosis Spinal Disord. 2017;12:14.

46. De Martinis M, Sirufo MM, Ginaldi L. Allergy and Aging: An Old/New Emerging Health Issue. Aging Dis. 2017;8(2):162–175.

47. Heymann D, Stern Y, Cosentino S, Tatarina-Nulman O, Dorrejo JN, Gu Y. The Association Between Alcohol Use and the Progression of Alzheimer’s Disease. Curr Alzheimer Res. 2016;13(12):1356–1362.

48. Durazzo TC, Mattsson N, Weiner MW, Alzheimer’s Disease Neuroimaging Initiative. Smoking and increased Alzheimer’s disease risk: a review of potential mechanisms. Alzheimers Dement. 2014;10(3 Suppl):S122–S145.

49. Dowling GJ, Weiss SRB, Condon TP. Drugs of abuse and the aging brain. Neuropsychopharmacology. 2008;33(2):209–218.

50. Arvanitakis Z, Capuano AW, Lamar M, et al. Late-life blood pressure association with cerebrovascular and Alzheimer disease pathology. Neurology. 2018;91(6):e517–e525.

51. Puglielli L, Tanzi RE, Kovacs DM. Alzheimer’s disease: the cholesterol connection. Nat Neurosci. 2003;6(4):345–351.

52. Huang ES, Laiteerapong N, Liu JY, John PM, Moffet HH, Karter AJ. Rates of complications and mortality in older patients with diabetes mellitus: the diabetes and aging study. JAMA Intern Med. 2014;174(2):251–258.

53. Marques JP, Kober T, Krueger G, van der Zwaag W, Van de Moortele PF, Gruetter R. MP2RAGE, a self bias-field corrected sequence for improved segmentation and T1-mapping at high field. Neuroimage. 2010;49(2):1271–1281.

54. Bedford SA, Park MTM, Devenyi GA, et al. Large-scale analyses of the relationship between sex, age and intelligence quotient heterogeneity and cortical morphometry in autism spectrum disorder. Mol Psychiatry. 2020;25(3):614–628.

55. Bellon EM, Haacke EM, Coleman PE, Sacco DC, Steiger DA, Gangarosa RE. MR artifacts: a review. AJR Am J Roentgenol. 1986;147(6):1271–1281.

56. Smith TB, Nayak KS. MRI artifacts and correction strategies. Imaging Med. Published online 2010. https://www.openaccessjournals.com/articles/mri-artifacts-and-correction-strategies-11010.html

57. Stüber C, Morawski M, Schäfer A, et al. Myelin and iron concentration in the human brain: A quantitative study of MRI contrast. Neuroimage. 2014;93:95–106.

58. Tustison NJ, Avants BB, Cook PA, et al. N4ITK: improved N3 bias correction. IEEE Trans Med Imaging. 2010;29(6):1310–1320.

59. Eskildsen SF, Coupé P, Fonov V, et al. BEaST: brain extraction based on nonlocal segmentation technique. Neuroimage. 2012;59(3):2362–2373.

60. Collins DL, Neelin P, Peters TM, Evans AC. Automatic 3D intersubject registration of MR volumetric data in standardized Talairach space. J Comput Assist Tomogr. 1994;18(2):192–205.

61. Kim JS, Singh V, Lee JK, et al. Automated 3-D extraction and evaluation of the inner and outer cortical surfaces using a Laplacian map and partial volume effect classification. Neuroimage. 2005;27(1):210–221.

62. Paquola C, Vos De Wael R, Wagstyl K, et al. Microstructural and functional gradients are increasingly dissociated in transmodal cortices. PLoS Biol. 2019;17(5):e3000284.

63. Whitaker KJ, Vértes PE, Romero-Garcia R, et al. Adolescence is associated with genomically patterned consolidation of the hubs of the human brain connectome. Proc Natl Acad Sci U S A. 2016;113(32):9105–9110.

64. Cohen-Adad J, Polimeni JR, Helmer KG, et al. T[* mapping and B[ orientation-dependence at 7 T reveal cyto- and myeloarchitecture organization of the human cortex. Neuroimage. 2012;60(2):1006–1014.

65. Chung MK, Adluru N, Vorperian HK. Heat Kernel Smoothing on Manifolds and Its Application to Hyoid Bone Growth Modeling. In: Zhao Y, Chen DG (din), eds. Statistical Modeling in Biomedical Research: Contemporary Topics and Voices in the Field. Springer International Publishing; 2020:235–261.

66. Avants BB, Tustison NJ, Song G, Cook PA, Klein A, Gee JC. A reproducible evaluation of ANTs similarity metric performance in brain image registration. Neuroimage. 2011;54(3):2033–2044.

67. Bussy A, Levy JP, Best T, et al. Cerebellar and subcortical atrophy contribute to psychiatric symptoms in frontotemporal dementia. Hum Brain Mapp. 2023;44(7):2684–2700.

68. Chung MK, Worsley KJ, Paus T, et al. A unified statistical approach to deformation-based morphometry. Neuroimage. 2001;14(3):595–606.

69. Sotiras A, Resnick SM, Davatzikos C. Finding imaging patterns of structural covariance via Non-Negative Matrix Factorization. Neuroimage. 2015;108:1–16.

70. Robert C, Patel R, Blostein N, Steele CC, Mallar Chakravarty M. Analyses of microstructural variation in the human striatum using non-negative matrix factorization. Neuroimage. Published online November 27, 2021:118744.

71. Patel R, Mackay CE, Jansen MG, et al. Inter- and intra-individual variation in brain structural-cognition relationships in aging. Neuroimage. 2022;257:119254.

72. Kalantar-Hormozi H, Patel R, Dai A, et al. A cross-sectional and longitudinal study of human brain development: the integration of cortical thickness, surface area, gyrification index, and cortical curvature into a unified analytical framework. Neuroimage. Published online 2023:119885.

73. Marek S, Tervo-Clemmens B, Calabro FJ, et al. Publisher Correction: Reproducible brain-wide association studies require thousands of individuals. Nature. 2022;605(7911):E11.

74. McIntosh AR, Lobaugh NJ. Partial least squares analysis of neuroimaging data: applications and advances. Neuroimage. 2004;23 Suppl 1:S250–S263.

75. Kovacevic N, Abdi H, Beaton D, McIntosh AR. Revisiting PLS Resampling: Comparing Significance Versus Reliability Across Range of Simulations. In: New Perspectives in Partial Least Squares and Related Methods. Springer New York; 2013:159–170.

76. Zeighami Y, Fereshtehnejad SM, Dadar M, et al. A clinical-anatomical signature of Parkinson’s disease identified with partial least squares and magnetic resonance imaging. Neuroimage. 2019;190:69–78.

77. Dickerson BC, Bakkour A, Salat DH, et al. The cortical signature of Alzheimer’s disease: regionally specific cortical thinning relates to symptom severity in very mild to mild AD dementia and is detectable in asymptomatic amyloid-positive individuals. Cereb Cortex. 2009;19(3):497–510.

78. Singh V, Chertkow H, Lerch JP, Evans AC, Dorr AE, Kabani NJ. Spatial patterns of cortical thinning in mild cognitive impairment and Alzheimer’s disease. Brain. 2006;129(Pt 11):2885–2893.

79. Iannopollo E, Garcia K, Alzheimer’ Disease Neuroimaging Initiative. Enhanced detection of cortical atrophy in Alzheimer’s disease using structural MRI with anatomically constrained longitudinal registration. Hum Brain Mapp. 2021;42(11):3576–3592.

80. Xiong RM, Xie T, Zhang H, et al. The pattern of cortical thickness underlying disruptive behaviors in Alzheimer’s disease. psychoradiology. 2022;2(3):113–120.

81. Telegina DV, Suvorov GK, Kozhevnikova OS, Kolosova NG. Mechanisms of Neuronal Death in the Cerebral Cortex during Aging and Development of Alzheimer’s Disease-Like Pathology in Rats. Int J Mol Sci. 2019;20(22). doi:10.3390/ijms20225632

82. Carmeli C, Donati A, Antille V, et al. Demyelination in mild cognitive impairment suggests progression path to Alzheimer’s disease. PLoS One. 2013;8(8):e72759.

83. Luo X, Li K, Zeng Q, et al. Application of T1-/T2-Weighted Ratio Mapping to Elucidate Intracortical Demyelination Process in the Alzheimer’s Disease Continuum. Front Neurosci. 2019;13:904.

84. Zhao Z, Zhang L, Wen Q, et al. The effect of beta-amyloid and tau protein aggregations on magnetic susceptibility of anterior hippocampal laminae in Alzheimer’s diseases. Neuroimage. 2021;244:118584.

85. Livingston G, Sommerlad A, Orgeta V, et al. Dementia prevention, intervention, and care. Lancet. 2017;390(10113):2673–2734.

86. Teri L, Ferretti LE, Gibbons LE, et al. Anxiety of Alzheimer’s disease: prevalence, and comorbidity. J Gerontol A Biol Sci Med Sci. 1999;54(7):M348–M352.

87. Liew TM. Subjective cognitive decline, anxiety symptoms, and the risk of mild cognitive impairment and dementia. Alzheimers Res Ther. 2020;12(1):107.

88. Porter VR, Buxton WG, Fairbanks LA, et al. Frequency and characteristics of anxiety among patients with Alzheimer’s disease and related dementias. J Neuropsychiatry Clin Neurosci. 2003;15(2):180–186.

89. Kaiser NC, Liang LJ, Melrose RJ, Wilkins SS, Sultzer DL, Mendez MF. Differences in anxiety among patients with early-versus late-onset Alzheimer’s disease. J Neuropsychiatry Clin Neurosci. 2014;26(1):73–80.

90. Trofimova O, Loued-Khenissi L, DiDomenicantonio G, et al. Brain tissue properties link cardio-vascular risk factors, mood and cognitive performance in the CoLaus|PsyCoLaus epidemiological cohort. Neurobiol Aging. 2021;102:50–63.

91. Pink A, Przybelski SA, Krell-Roesch J, et al. Cortical Thickness and Anxiety Symptoms Among Cognitively Normal Elderly Persons: The Mayo Clinic Study of Aging. J Neuropsychiatry Clin Neurosci. 2017;29(1):60–66.

92. George KM, Maillard P, Gilsanz P, et al. Association of Early Adulthood Hypertension and Blood Pressure Change With Late-Life Neuroimaging Biomarkers. JAMA Netw Open. 2023;6(4):e236431.

93. Gonzalez CE, Pacheco J, Beason-Held LL, Resnick SM. Longitudinal changes in cortical thinning associated with hypertension. J Hypertens. 2015;33(6):1242–1248.

94. Fungwe TV, Ngwa JS, Johnson SP, et al. Systolic Blood Pressure Is Associated with Increased Brain Amyloid Load in Mild Cognitively Impaired Participants: Alzheimer’s Disease Neuroimaging Initiatives Study. Dement Geriatr Cogn Disord. Published online February 20, 2023:1–8.

95. Skoog I, Gustafson D. Update on hypertension and Alzheimer’s disease. Neurol Res. 2006;28(6):605–611.

96. Sparks DL, Scheff SW, Hunsaker JC 3rd, Liu H, Landers T, Gross DR. Induction of Alzheimer-like beta-amyloid immunoreactivity in the brains of rabbits with dietary cholesterol. Exp Neurol. 1994;126(1):88–94.

97. Refolo LM, Malester B, LaFrancois J, et al. Hypercholesterolemia accelerates the Alzheimer’s amyloid pathology in a transgenic mouse model. Neurobiol Dis. 2000;7(4):321–331.

98. Umeda T, Tomiyama T, Kitajima E, et al. Hypercholesterolemia accelerates intraneuronal accumulation of Aβ oligomers resulting in memory impairment in Alzheimer’s disease model mice. Life Sci. 2012;91(23-24):1169–1176.

99. Refolo LM, Pappolla MA, LaFrancois J, et al. A cholesterol-lowering drug reduces beta-amyloid pathology in a transgenic mouse model of Alzheimer’s disease. Neurobiol Dis. 2001;8(5):890–899.

100. Olmastroni E, Molari G, De Beni N, et al. Statin use and risk of dementia or Alzheimer’s disease: a systematic review and meta-analysis of observational studies. Eur J Prev Cardiol. 2022;29(5):804–814.

101. Montagne A, Barnes SR, Sweeney MD, et al. Blood-brain barrier breakdown in the aging human hippocampus. Neuron. 2015;85(2):296–302.

102. Yanagisawa K. Cholesterol and amyloid beta fibrillogenesis. Subcell Biochem. 2005;38:179–202.

103. Popugaeva E, Pchitskaya E, Bezprozvanny I. Dysregulation of Intracellular Calcium Signaling in Alzheimer’s Disease. Antioxid Redox Signal. 2018;29(12):1176–1188.

104. Thelen KM, Falkai P, Bayer TA, Lütjohann D. Cholesterol synthesis rate in human hippocampus declines with aging. Neurosci Lett. 2006;403(1-2):15–19.

105. Blanchard JW, Akay LA, Davila-Velderrain J, et al. APOE4 impairs myelination via cholesterol dysregulation in oligodendrocytes. Nature. 2022;611(7937):769–779.

106. Vance JE. Dysregulation of cholesterol balance in the brain: contribution to neurodegenerative diseases. Dis Model Mech. 2012;5(6):746–755.

107. Hatters DM, Peters-Libeu CA, Weisgraber KH. Apolipoprotein E structure: insights into function. Trends Biochem Sci. 2006;31(8):445–454.

108. Chen R. Association of environmental tobacco smoke with dementia and Alzheimer’s disease among never smokers. Alzheimers Dement. 2012;8(6):590–595.

109. Cataldo JK, Prochaska JJ, Glantz SA. Cigarette smoking is a risk factor for Alzheimer’s Disease: an analysis controlling for tobacco industry affiliation. J Alzheimers Dis. 2010;19(2):465–480.

110. Kühn S, Schubert F, Gallinat J. Reduced thickness of medial orbitofrontal cortex in smokers. Biol Psychiatry. 2010;68(11):1061–1065.

111. Brody AL, Mandelkern MA, Jarvik ME, et al. Differences between smokers and nonsmokers in regional gray matter volumes and densities. Biol Psychiatry. 2004;55(1):77–84.

112. Duriez Q, Crivello F, Mazoyer B. Sex-related and tissue-specific effects of tobacco smoking on brain atrophy: assessment in a large longitudinal cohort of healthy elderly. Front Aging Neurosci. 2014;6:299.

113. Almeida OP, Garrido GJ, Lautenschlager NT, Hulse GK, Jamrozik K, Flicker L. Smoking is associated with reduced cortical regional gray matter density in brain regions associated with incipient Alzheimer disease. Am J Geriatr Psychiatry. 2008;16(1):92–98.

114. Tappia PS, Troughton KL. ṣcytokine production and antioxidant defences. Clin Sci. Published online 1995. https://europepmc.org/article/med/7540525

115. Bloomer RJ. Decreased blood antioxidant capacity and increased lipid peroxidation in young cigarette smokers compared to nonsmokers: impact of dietary intake. Nutr J. Published online 2007. https://nutritionj.biomedcentral.com/articles/10.1186/1475-2891-6-39

116. Lutti A, Dick F, Sereno MI, Weiskopf N. Using high-resolution quantitative mapping of R1 as an index of cortical myelination. Neuroimage. 2014;93:176–188.

117. Harkins KD, Xu J, Dula AN, et al. The microstructural correlates of T1 in white matter. Magn Reson Med. 2016;75(3):1341–1345.

118. Vymazal J, Brooks RA, Baumgarner C, et al. The relation between brain iron and NMR relaxation times: an in vitro study. Magn Reson Med. 1996;35(1):56–61.

119. Gelman N, Ewing JR, Gorell JM, Spickler EM, Solomon EG. Interregional variation of longitudinal relaxation rates in human brain at 3.0 T: relation to estimated iron and water contents. Magn Reson Med. 2001;45(1):71–79.

120. Miletić S, Bazin PL, Isherwood SJS, Keuken MC, Alkemade A, Forstmann BU. Charting human subcortical maturation across the adult lifespan with in vivo 7 T MRI. Neuroimage. 2022;249:118872x.

121. Brown RW, Cheng YCN, Mark Haacke E, Thompson MR, Venkatesan R. Magnetic Resonance Imaging: Physical Principles and Sequence Design. John Wiley & Sons; 2014.

122. Duyn J. MR susceptibility imaging. J Magn Reson. 2013;229:198–207.

123. Langkammer C, Krebs N, Goessler W, et al. Quantitative MR imaging of brain iron: a postmortem validation study. Radiology. 2010;257(2):455–462.

124. van der Weijden CWJ, García DV, Borra RJH, et al. Myelin quantification with MRI: A systematic review of accuracy and reproducibility. Neuroimage. 2021;226:117561.

125. Tardif CL, Bedell BJ, Eskildsen SF, Collins DL, Pike GB. Quantitative magnetic resonance imaging of cortical multiple sclerosis pathology. Mult Scler Int. 2012;2012:742018.

126. Schmierer K, Scaravilli F, Altmann DR, Barker GJ, Miller DH. Magnetization transfer ratio and myelin in postmortem multiple sclerosis brain. Ann Neurol. 2004;56(3):407–415.

127. Schmierer K, Wheeler-Kingshott CAM, Tozer DJ, et al. Quantitative magnetic resonance of postmortem multiple sclerosis brain before and after fixation. Magn Reson Med. 2008;59(2):268–277.

128. Odrobina EE, Lam TYJ, Pun T, Midha R, Stanisz GJ. MR properties of excised neural tissue following experimentally induced demyelination. NMR Biomed. 2005;18(5):277–284.

129. Thiessen JD, Zhang Y, Zhang H, et al. Quantitative MRI and ultrastructural examination of the cuprizone mouse model of demyelination. NMR Biomed. 2013;26(11):1562–1581.

130. Wood JC, Ghugre N. Magnetic resonance imaging assessment of excess iron in thalassemia, sickle cell disease and other iron overload diseases. Hemoglobin. 2008;32(1-2):85–96.

131. Hernando D, Levin YS, Sirlin CB, Reeder SB. Quantification of liver iron with MRI: state of the art and remaining challenges. J Magn Reson Imaging. 2014;40(5):1003–1021.

132. Walsh AJ, Lebel RM, Eissa A, et al. Multiple sclerosis: validation of MR imaging for quantification and detection of iron. Radiology. 2013;267(2):531–542.

133. Bagnato F, Hametner S, Boyd E, et al. Untangling the R2* contrast in multiple sclerosis: A combined MRI-histology study at 7.0 Tesla. PLoS One. 2018;13(3):e0193839.

134. Birkl C, Birkl-Toeglhofer AM, Kames C, et al. The influence of iron oxidation state on quantitative MRI parameters in post mortem human brain. Neuroimage. 2020;220:117080.

135. Kirsch SJ, Jacobs RW, Butcher LL, Beatty J. Prolongation of magnetic resonance T2 time in hippocampus of human patients marks the presence and severity of Alzheimer’s disease. Neurosci Lett. 1992;134(2):187–190.

136. Laakso MP, Partanen K, Soininen H, et al. MR T2 relaxometry in Alzheimer’s disease and age-associated memory impairment. Neurobiol Aging. 1996;17(4):535–540.

137. Wang H, Yuan H, Shu L, Xie J, Zhang D. Prolongation of T(2) relaxation times of hippocampus and amygdala in Alzheimer’s disease. Neurosci Lett. 2004;363(2):150–153.

138. Luo Z, Zhuang X, Kumar D, et al. The correlation of hippocampal T2-mapping with neuropsychology test in patients with Alzheimer’s disease. PLoS One. 2013;8(9):e76203.

139. Campeau NG, Petersen RC, Felmlee JP, O’Brien PC, Jack CR Jr. Hippocampal transverse relaxation times in patients with Alzheimer disease. Radiology. 1997;205(1):197–201.

140. Wearn AR, Nurdal V, Saunders-Jennings E, et al. T2 heterogeneity: a novel marker of microstructural integrity associated with cognitive decline in people with mild cognitive impairment. Alzheimers Res Ther. 2020;12(1):105.

141. Reinert A, Morawski M, Seeger J, Arendt T, Reinert T. Iron concentrations in neurons and glial cells with estimates on ferritin concentrations. BMC Neurosci. 2019;20(1):25.

142. Filo S, Shaharabani R, Bar Hanin D, et al. Non-invasive assessment of normal and impaired iron homeostasis in the brain. Nat Commun. 2023;14(1):5467.

143. Walker LC, Jucker M. The Exceptional Vulnerability of Humans to Alzheimer’s Disease. Trends Mol Med. 2017;23(6):534–545.

144. Ackley SF, Zimmerman SC, Brenowitz WD, et al. Effect of reductions in amyloid levels on cognitive change in randomized trials: instrumental variable meta-analysis. BMJ. 2021;372:n156.

145. Bennett DA, Schneider JA, Arvanitakis Z, et al. Neuropathology of older persons without cognitive impairment from two community-based studies. Neurology. 2006;66(12):1837–1844.

146. Herrup K. The case for rejecting the amyloid cascade hypothesis. Nat Neurosci. 2015;18(6):794–799.

147. Kametani F, Hasegawa M. Reconsideration of Amyloid Hypothesis and Tau Hypothesis in Alzheimer’s Disease. Front Neurosci. 2018;12:25.

148. Braak H, Braak E. Development of Alzheimer-related neurofibrillary changes in the neocortex inversely recapitulates cortical myelogenesis. Acta Neuropathol. 1996;92(2):197–201.

149. Bartzokis G. Age-related myelin breakdown: a developmental model of cognitive decline and Alzheimer’s disease. Neurobiol Aging. 2004;25(1):5–18; author reply 49-62.

150. Pakkenberg B, Pelvig D, Marner L, et al. Aging and the human neocortex. Exp Gerontol. 2003;38(1-2):95–99.

151. Rao YL, Ganaraja B, Murlimanju BV, Joy T, Krishnamurthy A, Agrawal A. Hippocampus and its involvement in Alzheimer’s disease: a review. 3 Biotech. 2022;12(2):55.

152. Boudreau M, Tardif CL, Stikov N, Sled JG, Lee W, Pike GB. B1 mapping for bias-correction in quantitative T 1 imaging of the brain at 3T using standard pulse sequences. J Magn Reson Imaging. 2017;46(6):1673–1682.

153. Bender B, Klose U. The in vivo influence of white matter fiber orientation towards B(0) on T2* in the human brain. NMR Biomed. 2010;23(9):1071–1076.

154. Melhem ER, Mori S, Mukundan G, Kraut MA, Pomper MG, van Zijl PCM. Diffusion tensor MR imaging of the brain and white matter tractography. AJR Am J Roentgenol. 2002;178(1):3–16.

155. Tax CMW, Kleban E, Chamberland M, Baraković M, Rudrapatna U, Jones DK. Measuring compartmental T2-orientational dependence in human brain white matter using a tiltable RF coil and diffusion-T2 correlation MRI. Neuroimage. 2021;236:117967.

