## Supplementary material for "Joint signatures of morphological and microstructural inter-individual variation in the Alzheimer’s spectrum"

1. **Participant’s inclusion and exclusion**

Individuals were recruited across two distinct studies: the Alzheimer's Disease Biomarkers (ADB) Study and the PREVENT-AD Study. All participants provided signed informed consent and the study received approval from the Research Ethics Board of the Douglas Mental Health University Institute, Montreal, QC, Canada.

**ADB study**

Inclusion criteria for all participants in the study are as follows:

- Competency to provide informed consent (refer to section 6.5 on page 13).
- Age 55 and older.
- Fluency in English or French.

Exclusion criteria for all participants include:

- Comorbidity with neurological diseases unrelated to Mild Cognitive Impairment (MCI) and Alzheimer's Disease (AD) for MCI and AD cohorts.
- Comorbidity with axis I psychiatric disorders.
- Major structural neurological abnormalities or pathologies unrelated to MCI and AD for MCI and AD cohorts, including a history of brain damage or concussion.
- Current use of psychoactive substances, including a history of substance abuse or drug addiction, and ongoing or recent (within three months) treatment with medication known to affect serotonin neurotransmission.
- Intellectual disability, including a history of cognitive deficits unrelated to MCI or AD (e.g., learning disabilities).
- Less than 6 years of formal education.
- Possession of any contraindication to MRI, such as cardiac pacemaker, defibrillator, cochlear implant, or other metallic objects in the body, cardiac valve prosthesis, aneurysm clip, neurostimulators, claustrophobia, certain tattoos, filters, catheters, stents in blood vessels, or shunts, copper IUD, pregnancy, or other conditions preventing the participant from lying on their back or remaining still during the scan.

**Diagnostic criteria for MCI involve:**

- Age over 80 years.
- Presence of subjective memory complaints.
- Performance above the cutoff (24/30) for dementia in the Mini-Mental State Examination (MMSE).
- Performance above the cutoff for dementia on the Mattis Dementia Rating Scale.
- Clinical Dementia Rating (CDR) less than 0.5.
- Does not meet diagnostic criteria for probable or possible AD.
- Memory deficit higher than 1.5 standard deviations on a 3-delayed verbal recall test.
- No cognitive deficit higher than 1.5 standard deviations in other cognitive domains.

**Diagnostic criteria for mild AD include:**

- Meets DSM-IV criteria upon clinical assessment.
- Performance below the cutoff (18-24) for dementia in the MMSE.
- CDR=1.

**PREVENT-AD**

Participants adhered to inclusion criteria outlined by Tremblay-Mercier et al. (2021), including a self-reported parental or multiple-sibling history of Alzheimer-like dementia, age 60 or older, a minimum of 6 years of formal education, and proficiency in spoken and written French and/or English. Additional criteria involved having a study partner, fluency, intention to participate in regular visits, periodic blood and urine sample donations, and agreement to various assessments including MRI and lumbar puncture for cerebrospinal fluid collection.

Exclusion criteria were set for cognitive disorders identified during assessments, use of certain medications, hypertension, anemia, liver/kidney disease, and various substances. The comprehensive set of criteria ensured the study's robustness and the eligibility of participants.

2. **Cognitive, psychological, medical and lifestyle information**

The MOCA test has a score range of 0-30: a score of 26-30 is considered normal, 18-25 is indicative of mild cognitive impairment, 10-17 is indicative of moderate cognitive impairment, and a score below 10 is indicative of severe cognitive impairment. The RBANS test assesses multiple cognitive domains, including total score, immediate memory, delayed memory, attention, language, and visuospatial ability.

3. **Data imputation**

**Supplementary table 1:** **Table of the number of missing values for each of the variables per group.**

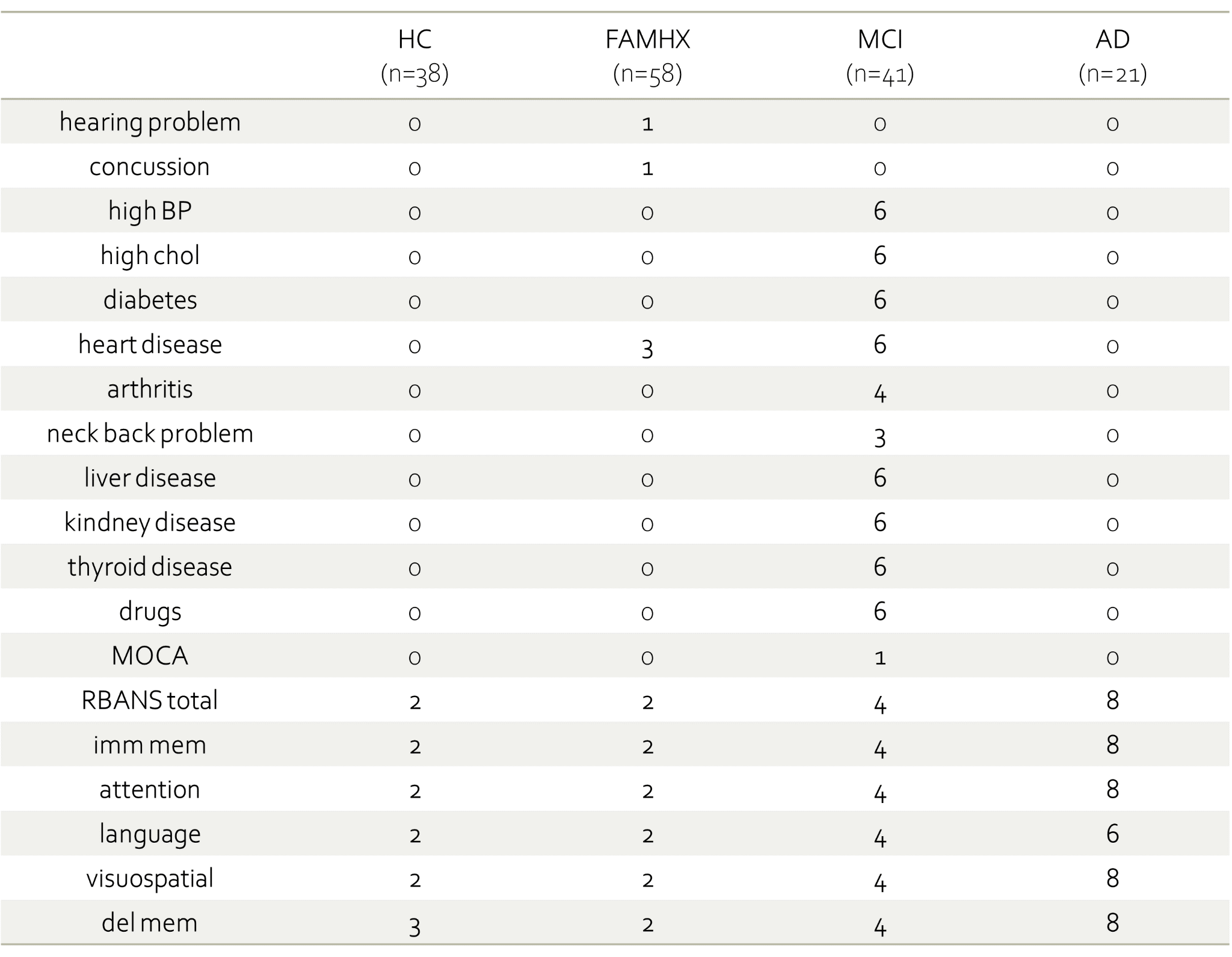

4. **T2* denoising**

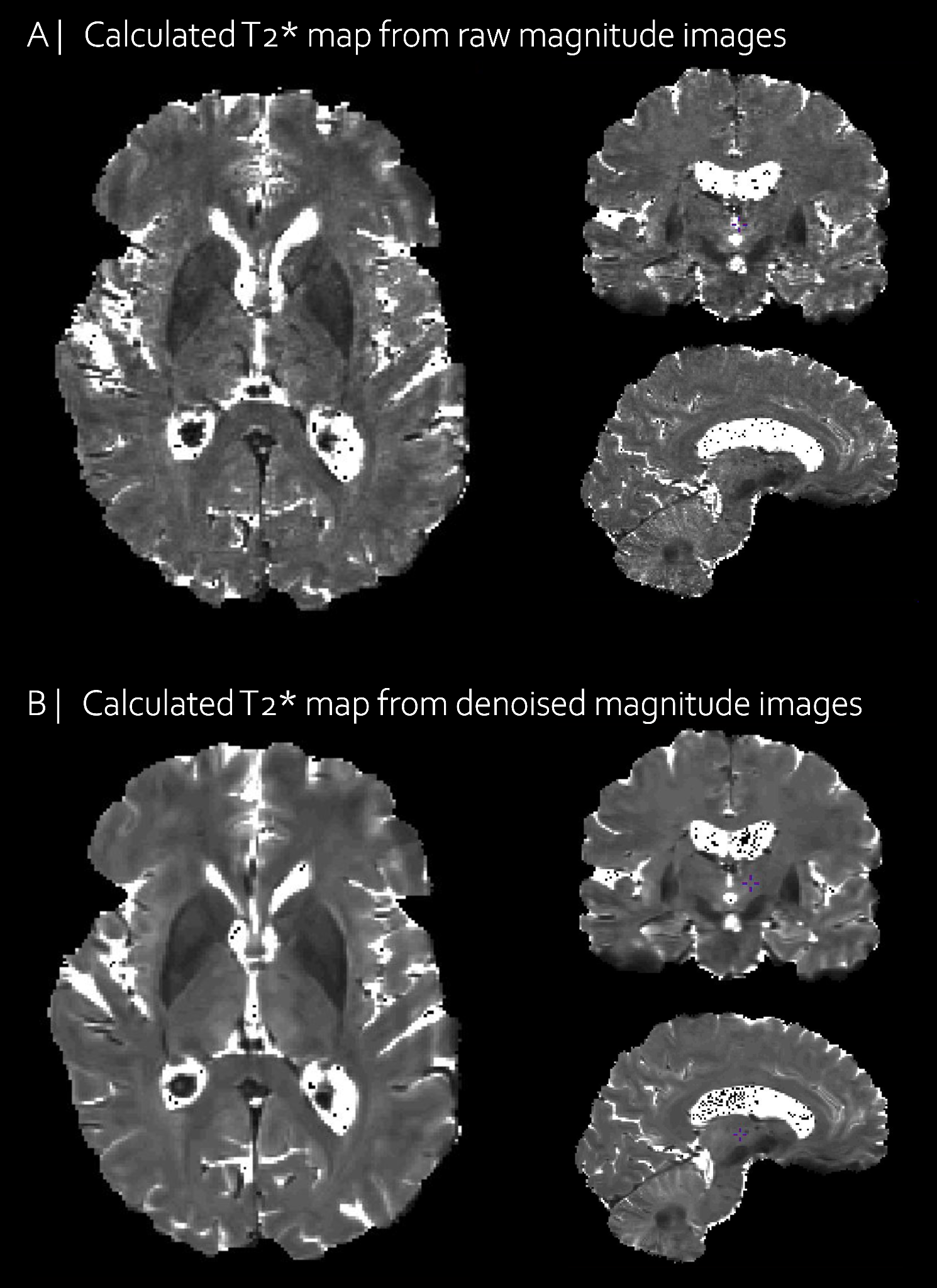

**Supplementary figure 1 :** T2* map example with or without denoising.

5. **T2* quality control (QC)**

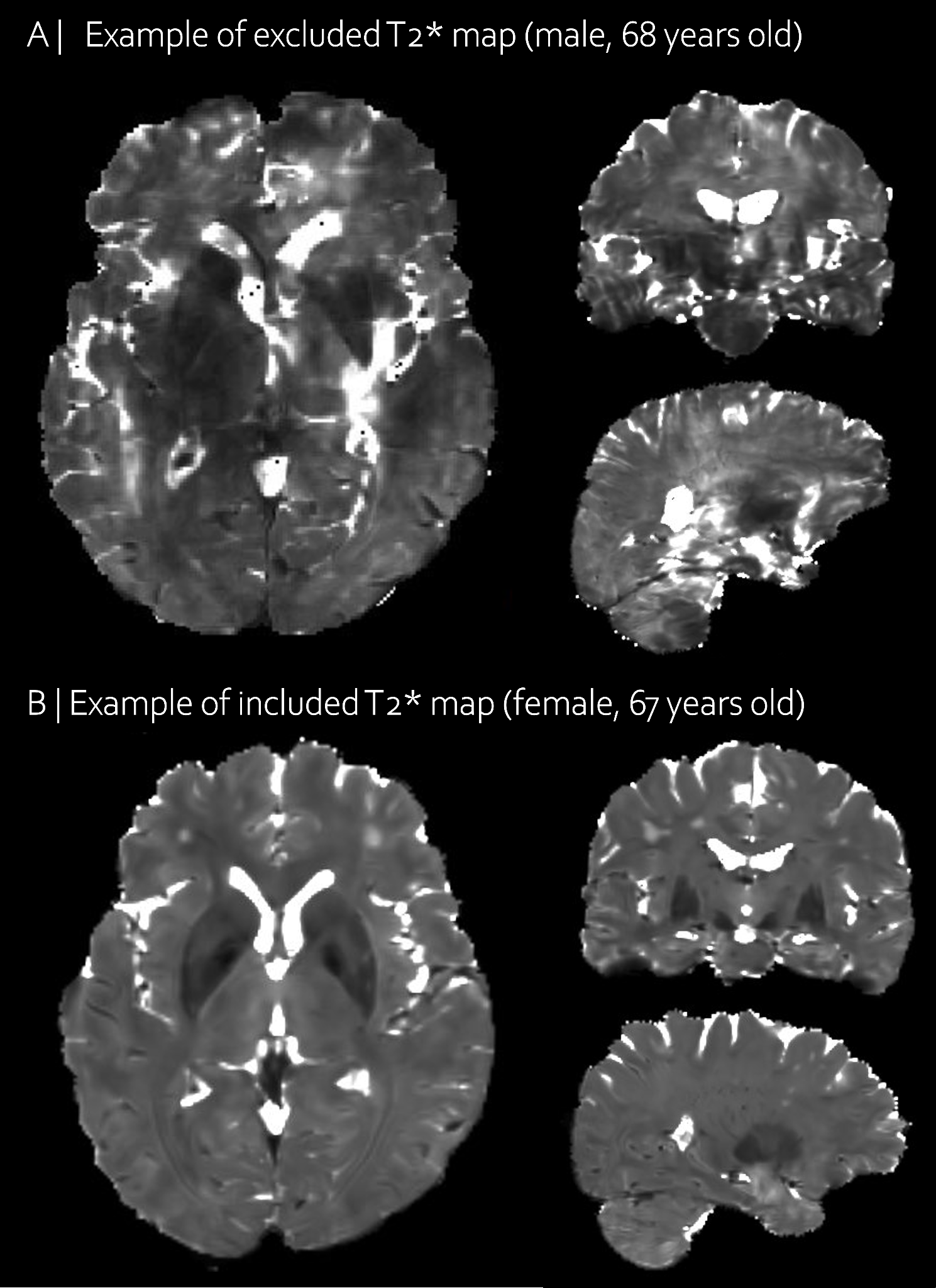

**Supplementary figure 2 :** A) T2* map example of an excluded participant due to motion artifacts and B) T2* map example of an included participant with good QC.

6. **Morphometric and qMRI cortical feature extraction**

To sample the maps vertex-wise, we first generated additional surfaces across multiple depths from the pial surface (“0% surface”) to the gray matter (GM) and white matter (WM) interface (“100% surface”) following similar approaches described by others(Paquola et al. 2019; Whitaker et al. 2016). Here we used increments of 12.5%, where each surface was generated by the vertex coordinates of two surfaces (e.g: 25% defined by averaging the coordinates of the 0% and 50% depths). T1 and T2* maps were sampled in their native space. Cortical surfaces were already in T1 space whereas for T2*, we first registered T1w images to the first echo of the T2* magnitude images and applied the same transformations to the cortical surfaces to have them in T2* space.

7. **Hippocampal mask creation**

MAGeT brain segmentation algorithm was used to segment the hippocampus of our participants. This technique has already been used and validated in our laboratory to extract hippocampal subfield labels, volumes and shape (Pipitone et al. 2014; Voineskos et al. 2015; Winterburn et al. 2013; Bussy et al. 2021; Bussy et al. 2021). After obtaining the hippocampal GM labels in native space, we transformed them into template space. A final hippocampal mask was obtained by performing a majority vote. Manual correction was used to improve this mask and to avoid voxels not labeled within the hippocampal structure (see **Supplementary figure 3**).

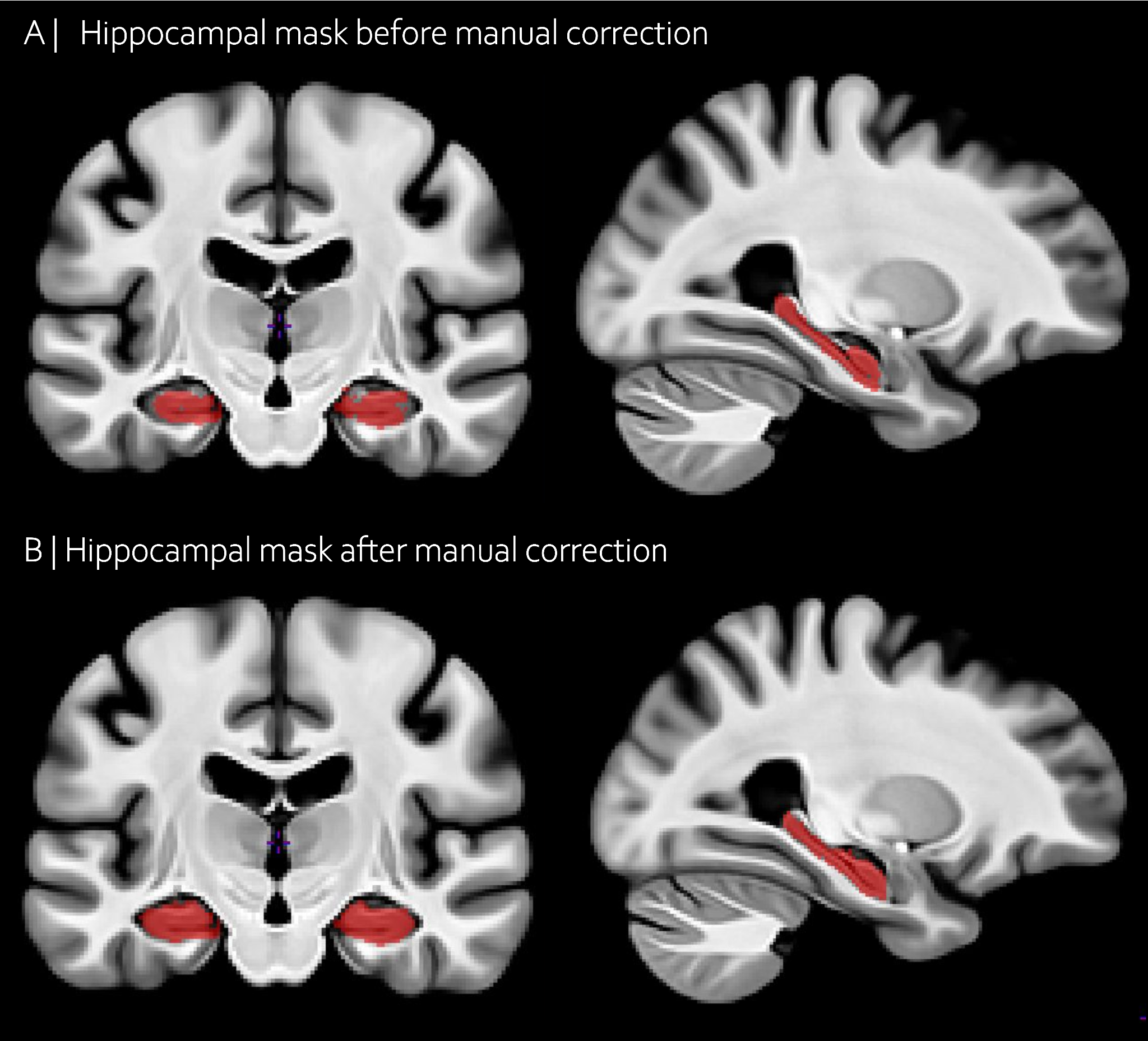

**Supplementary figure 3:** Hippocampal mask obtained after MAGeT brain **(A | )** Before manual correction and (**B | )** After manual correction. Red : voxels included in the mask.

8. **Description of NMF**

Here, we used orthogonal projective NMF to analyze cortical data (CT, SA, T1, T2*) and hippocampal data (J, T1, T2*)^17^. NMF decomposes a $m \times n$ input matrix into a component matrix $W (m\times k)$ and weight matrix $H (k\times n)$ constructed such that their multiplication minimizes the reconstruction error between the original and reconstructed data. The number of components $k$ is user defined. $W$ describes the component spatial locations, while $H$ contains subject-wise weightings describing individual variability of measures in each component. In the specific implementation used in this work, each column of the input matrix contains hemispheric cortical vertex-wise metrics $m=38,561$ or bilateral hippocampal voxel-wise $m=6,222$ and each row represents subject metrics ($number of rows=\#metrics \times158 subjects)$.

For the cortex, each hemispheric metric was initially concatenated to obtain one whole brain matrix $(77,122\times158$). This whole brain matrix was z-scored within each individual metric across vertices for the cortex (or voxels for the hippocampus) and participants **(Supplementary Figure 4**). NMF was run separately for the cortex and the hippocampus because of the difference in matrix size. Z-scored matrices were concatenated and the values were shifted by adding the minimal z-scored value across the 4 (or 3) matrices to have the minimum value equal to 0 (More details in **Supplementary Figure** **4)**.

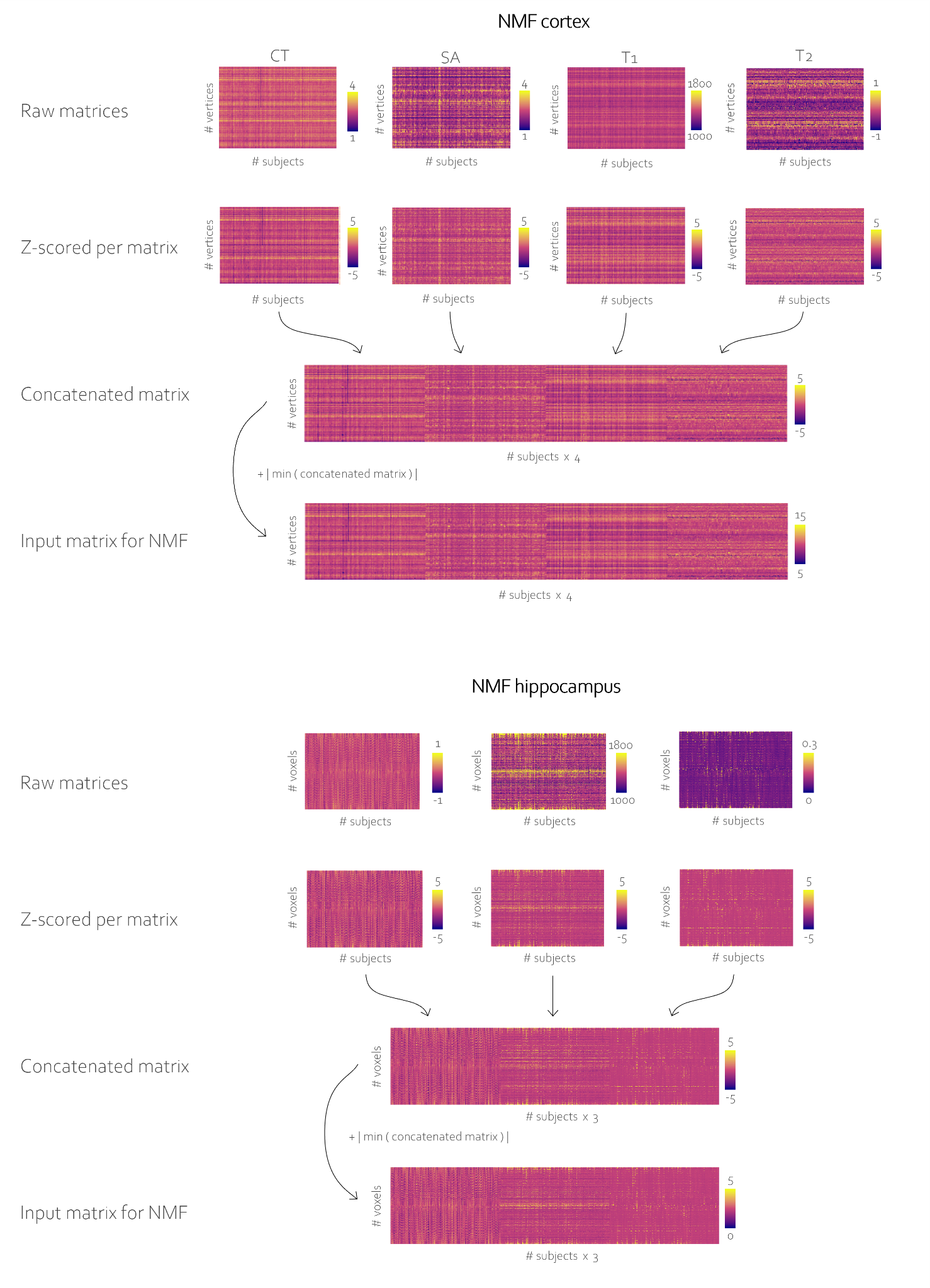

**Supplementary figure 4 : Cortical and hippocampal input matrix creation. I**nput matrices of the (**A |** ) vertex-wise cortical metrics and (**B |** ) voxel-wise hippocampal metrics. First, we performed a z-scoring across each input matrix (for example, across participants and vertices for CT). Then, the z-scored matrices were concatenated to create a large matrix (*number of columns = number of participants x number of metrics*). Finally, the absolute value of the minimum z-scored value across the large input matrix was added to all values to obtain a non-negative matrix (Patel et al. 2020).

9. **NMF stability analyses**

To select the optimal number of components, we performed a split half analysis from $k=2$ to $k=20$ to assess stability and accuracy of decompositions across different granularities^17^. Results of the stability analyses are in **Supplementary Figure 5**.

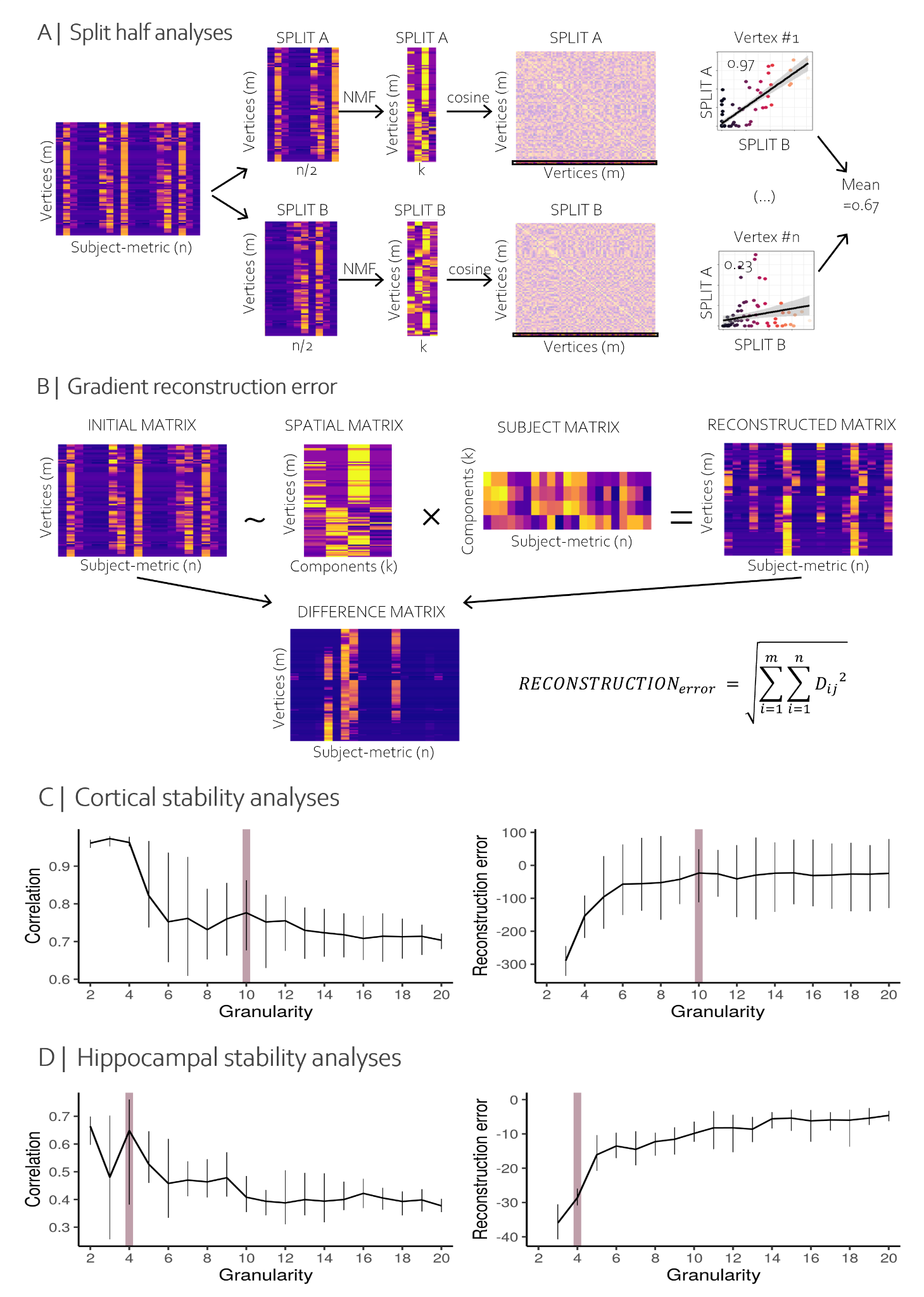

**Supplementary figure 5: Stability analyses results.** (**A |** ) Representation of the calculation of the correlation calculation between split A and split B. (**B |** ) Representation of the calculation of the gradient reconstruction error. (**C |** ) For the cortex, we decided to select 10 components to have a detailed enough parcellation, with reasonable correlation and reconstruction error. (**D|** ) For the hippocampus, we selected 4 components to have a high correlation and components of a decent size (also, previous work from our lab already selected 4 hippocampal components with a similar approach).

10. **Cortical NMF k4**

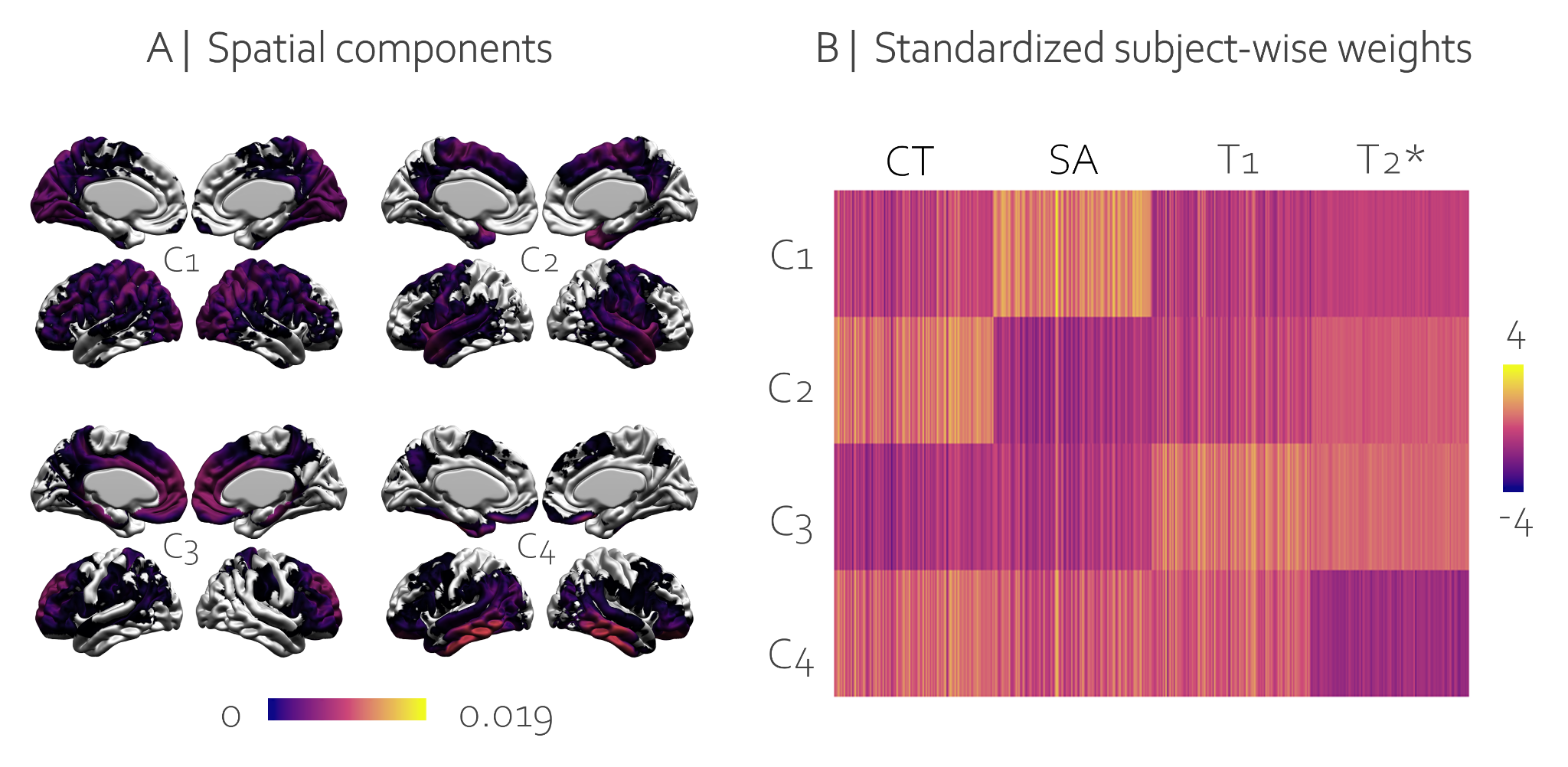

**Supplementary figure 6: Outputs for 4 components cortical NMF run.** (**A |** ) Spatial maps and (**B |** ) Standardized subject-wise weights matrix.

11. **Component characteristics**

Because the NMF output matrices are normalized per component, direct comparison of across components is not possible. Therefore, we used the NMF parcellation to extract the mean raw values to describe the components (**Supplementary Figure 7**).

The properties of orthogonal projection NMF enable us to assign each vertex (or voxel), via a winner takes all approach, such that each vertex (or voxel) was assigned to a single component for which it had the highest component score. Mean vertex-wise (or voxel-wise) values were calculated for each component and for each individual to visualize the raw value variation across the structures of interest and individuals to better characterize the variation captured by the components (**Supplementary Figure 7**). By doing so, we can appreciate the spatial variation of our metrics across our structures of interest, and we can visualize how variable the metrics are per component.

11.1. **Cortical components**

CT is higher in component 1, 4 and 5 which mostly corresponds to regions of the frontal, temporal and dorsomedial regions, while lower CT is seen in the occipital and sensorimotor regions. SA is mainly higher in the occipital lobe compared to the rest of the brain. T1 is higher in the frontal and temporal regions while T2* shows a gradient from high values in the occipital lobe to lower values in the occipital regions.

11.2. **Hippocampal components**

The distribution of J across participants is consistent in the four hippocampal components. However, higher T1 values were observed in the most lateral and the most medial regions of the hippocampus. T2* was higher in the lateral regions of the hippocampus.

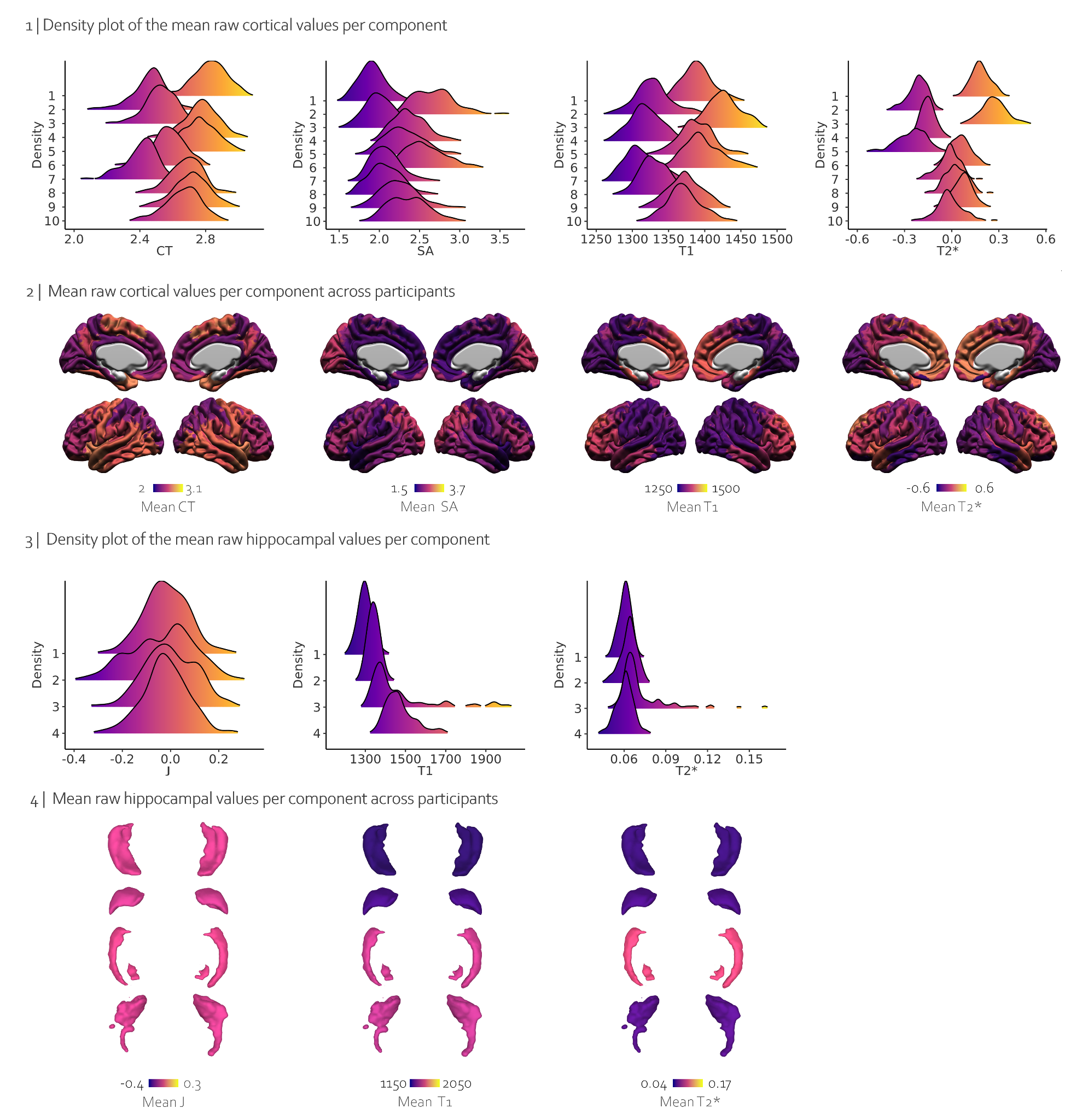

**Supplementary figure 7: Mean raw cortical and hippocampal values per component.** **1 |** Density plot of the mean raw cortical values per component. **2|** Mean raw cortical values per component across participants. **3 |** Density plot of the mean raw hippocampal values per component. **4 |** Mean raw hippocampal values per component across participants

12. **Statistical results of post-hoc analyses**

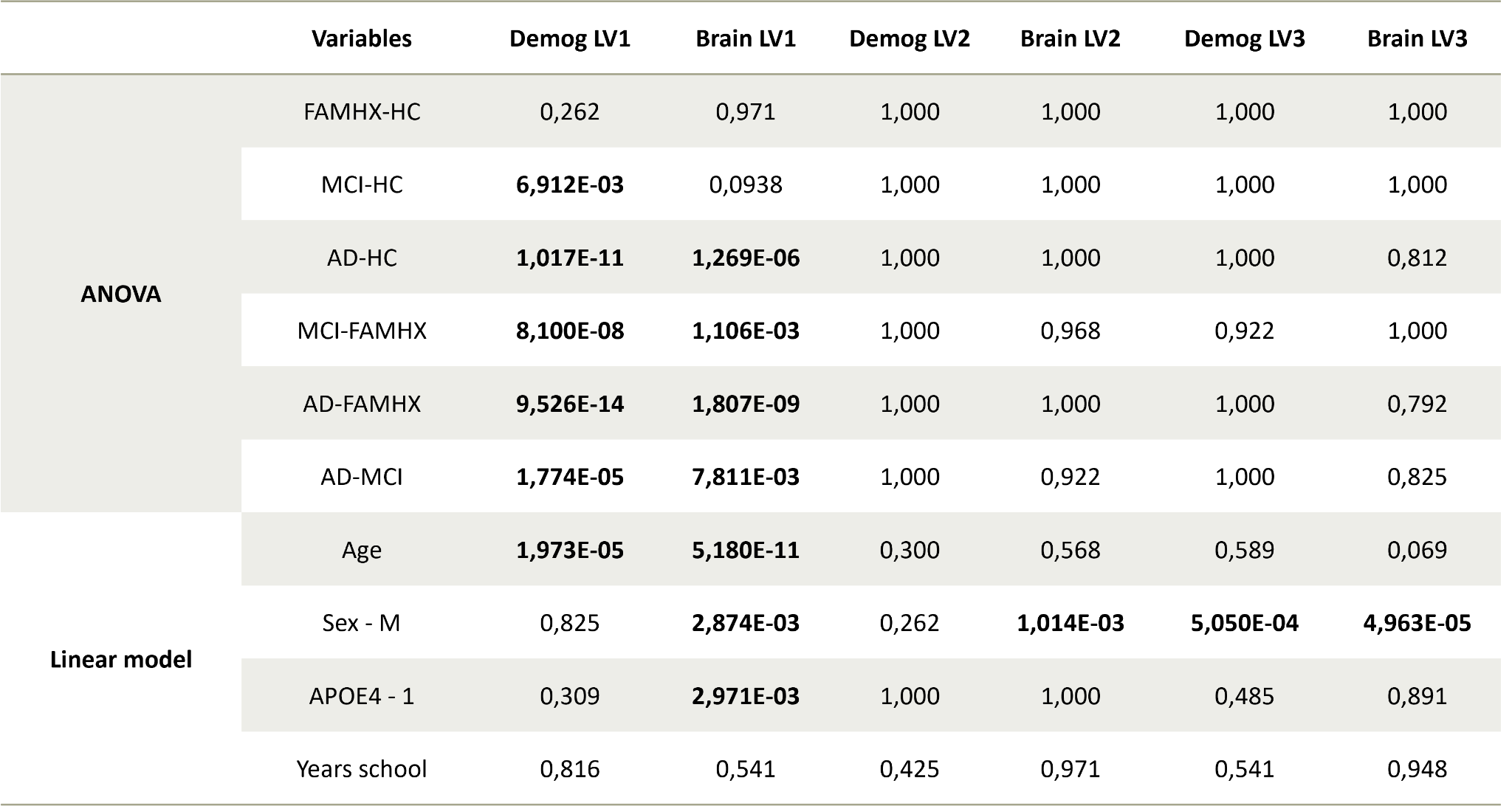

**Supplementary table 2 : Table of the q-values (p-values after FDR correction) across all variables and LVs.** Pairwise group differences were tested using ANOVA and post-hoc Tukey HSD, while age, sex, APOE4 and education were tested using linear models.

13. **LV2 results**

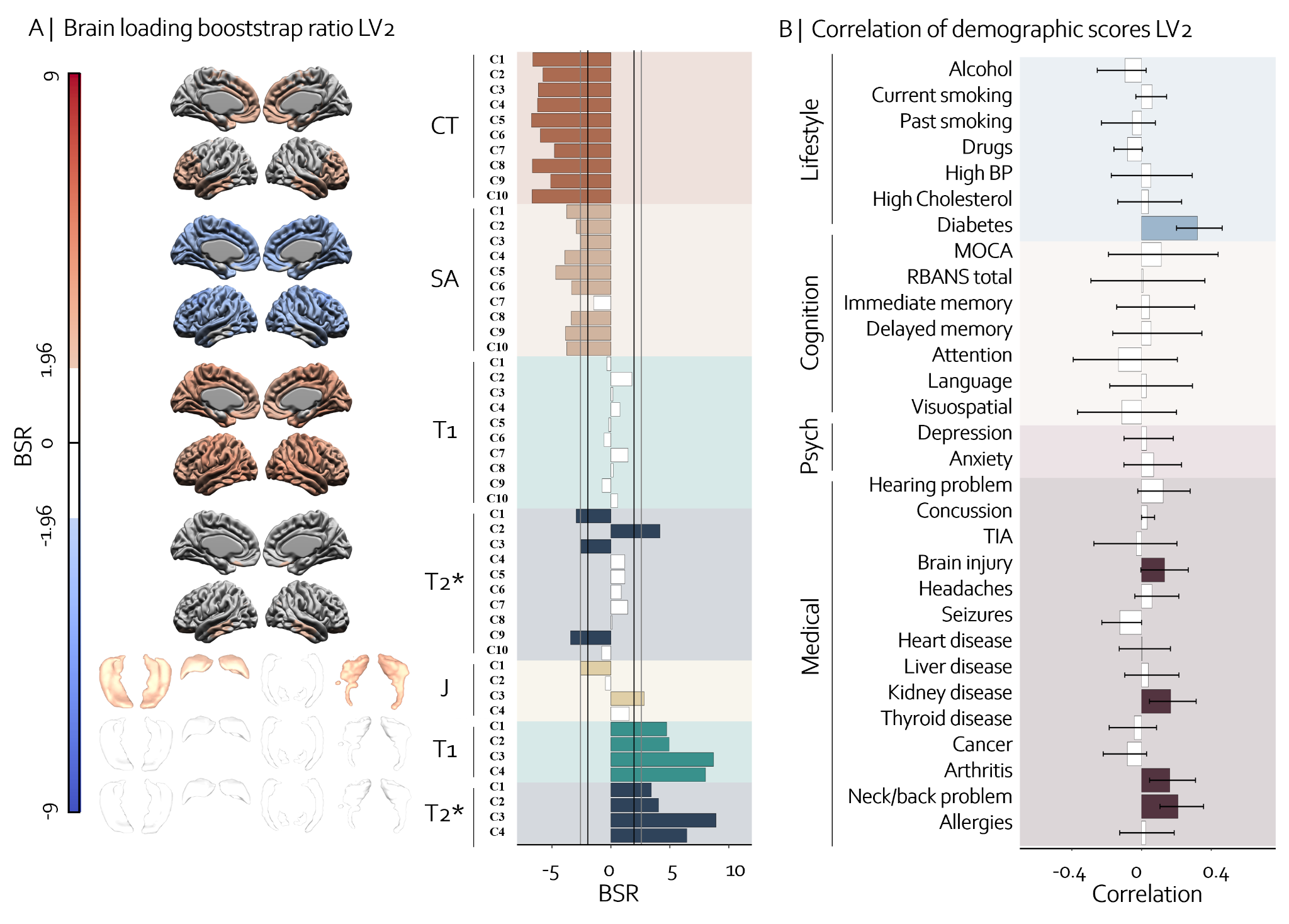
**Supplementary figure 8: Relationship between brain and cognitive, psychological, medical and lifestyle information of the LV2.** The second LV explained 7.5% of the covariance between brain and cognitive, psychological, medical, and lifestyle information. The brain pattern on the left **(A)** illustrates spatially the contribution of each metric to the pattern with BSR values on each brain structure. Blue color indicates negative BSR values and red indicates positive BSR values. Components with absolute BSR values higher than 1.96 are colored to show significant contribution. The bar plot in the middle of the figure shows more precisely the BSR values for each component, with black vertical lines representing a BSR of 1.96 (equivalent to p=0.05) and gray lines representing a BSR of 2.58 (equivalent to p=0.01). On the right-hand side **(B)**, we can see the cognitive, psychological, medical, and lifestyle patterns associated with the brain pattern on the left. Bars are colored if they are significant (when error bars do not cross zero), and are white if non-significant. The brain pattern is associated with having diabetes, brain injury, kidney disease, arthritis and neck/back problems.

14. **Post-hoc analyses of LV2**

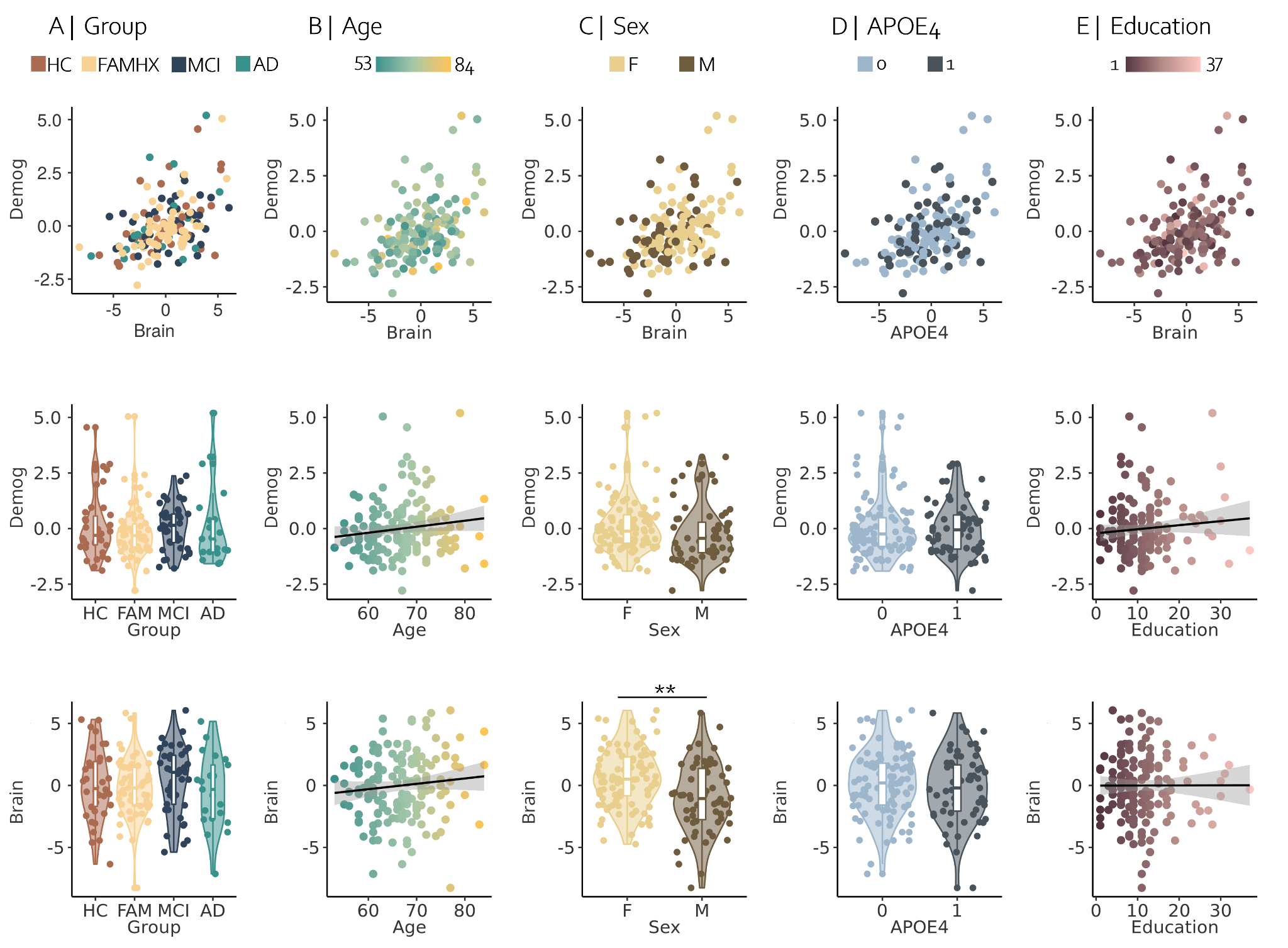

**Supplementary figure 9: Post-hoc analyses of the PLS brain and demographic scores of LV2. (A)** is showing the pairwise group comparisons of the brain and demographic scores. Results of the linear models illustrating the brain and demographic relationships with age **(B)**, sex **(C)**, APOE4 status **(D)** and education **(E)**. ***** p < 0.05, ****** p < 0.01, ******* p < 0.001 after FDR correction across all p-values.

15. **LV3 results**

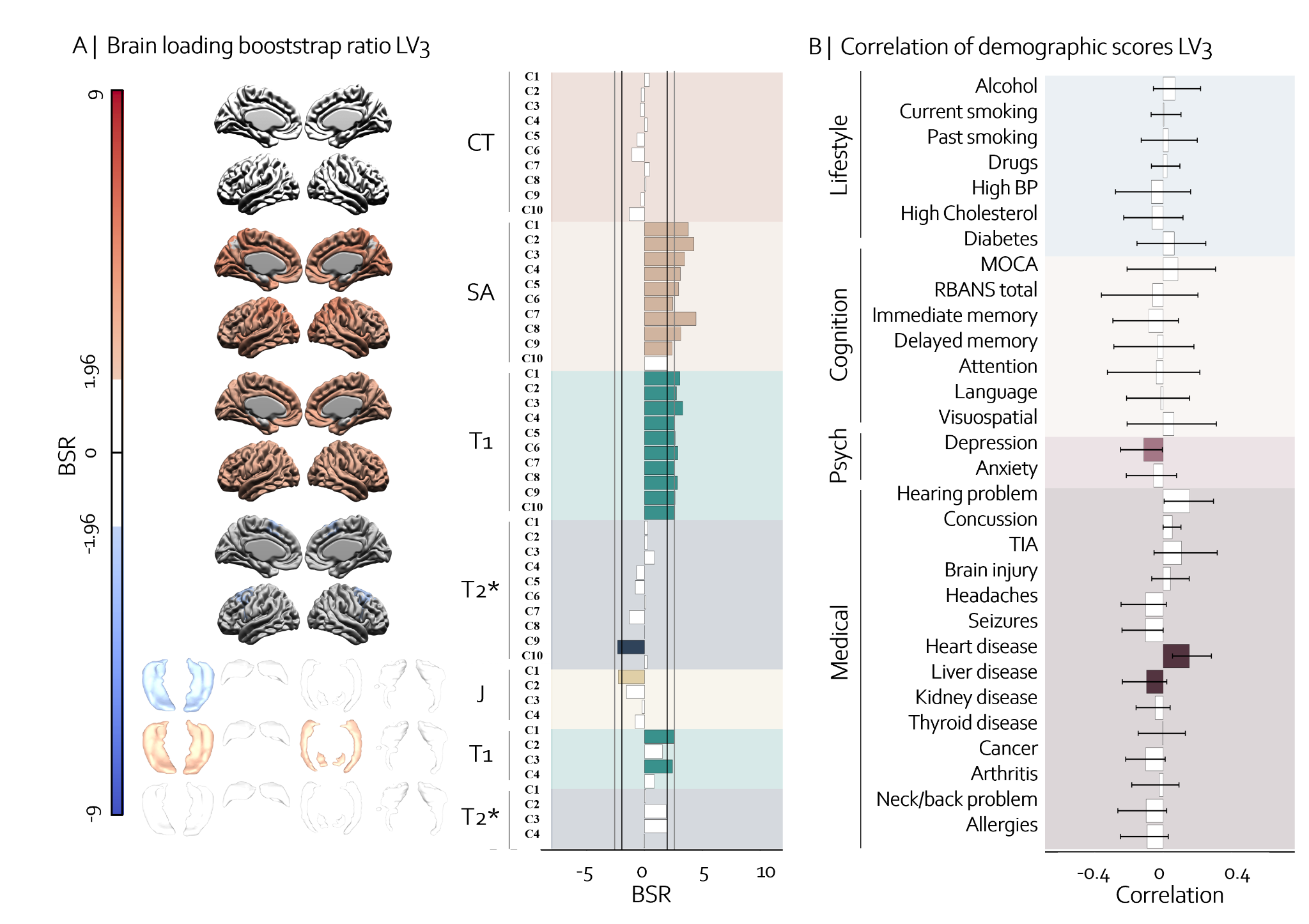
**Supplementary figure 10:** **Relationship between brain and cognitive, psychological, medical and lifestyle information of the LV3.** The third LV explained 5.2% of the covariance between brain and cognitive, psychological, medical, and lifestyle information. The brain pattern on the left **(A)** illustrates spatially the contribution of each metric to the pattern with BSR values on each brain structure. Blue color indicates negative BSR values and red indicates positive BSR values. Components with absolute BSR values higher than 1.96 are colored to show significant contribution. The bar plot in the middle of the figure shows more precisely the BSR values for each component, with black vertical lines representing a BSR of 1.96 (equivalent to p=0.05) and gray lines representing a BSR of 2.58 (equivalent to p=0.01). On the right-hand side **(B)**, we can see the cognitive, psychological, medical, and lifestyle patterns associated with the brain pattern on the left. Bars are colored if they are significant (when error bars do not cross zero), and are white if non-significant. The brain pattern is associated with having diabetes, brain injury, kidney disease, arthritis and neck/back problems.

16. **Post-hoc analyses of LV3**

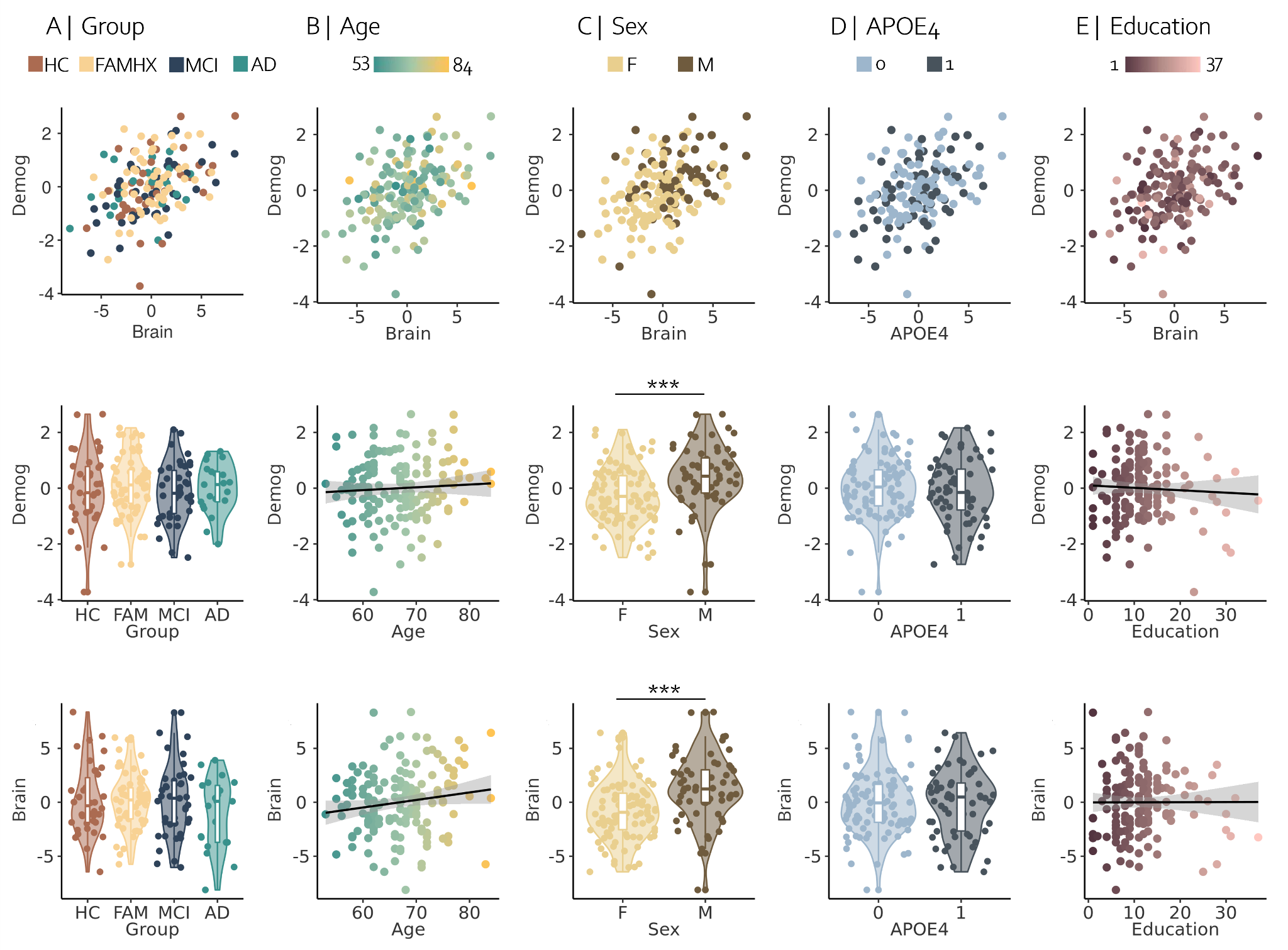

**Supplementary figure 11:** **Post-hoc analyses of the PLS brain and demographic scores of LV3. (A)** is showing the pairwise group comparisons of the brain and demographic scores. Results of the linear models illustrating the brain and demographic relationships with age **(B)**, sex **(C)**, APOE4 status **(D)** and education **(E)**. ***** p < 0.05, ****** p < 0.01, ******* p < 0.001 after FDR correction across all p-values.
